# Cancer-associated mutations reconfigure dynamical responses in EGFR kinase

**DOI:** 10.64898/2026.10.08.757680

**Authors:** Lorenzo Tulli, Alessio Lodola, Adrian J. Mulholland, A. Sofia F. Oliveira

## Abstract

The epidermal growth factor receptor (EGFR) is a tyrosine kinase receptor that plays a fundamental role in regulating cellular proliferation, survival, and differentiation. EGFR’s kinase domain catalyses the autophosphorylation that drives downstream signalling and is a known hotspot for cancer-associated mutations. Such substitutions often enhance receptor activation, driving oncogenesis and therapeutic resistance, with major implications for prognosis and treatment choice; consequently, a detailed understanding of this domain’s functional dynamics, and how mutations reshape them, is essential. Here, we apply dynamical-nonequilibrium molecular dynamics (D-NEMD) simulations to probe how changes at the ATP-binding site are transmitted throughout the kinase domain, ultimately shaping the behaviour of distal regions in the wild-type protein and in two of the most clinically observed non-small cell lung cancer variants, L858R and L858R+T790M. D-NEMD reveals previously uncharacterised communication networks linking the ATP pocket to key functional motifs, including the P-loop, αC-helix, hinge region, activation loop, and αF-helix, showing the intricate allosteric connectivity within this domain. Notably, numerous cancer-related mutation sites, despite being distant from the ATP site, map onto or lie close to these communication pathways, indicating that D-NEMD simulations can identify functionally relevant allosteric positions and potential mutation sites in EGFR. Our simulations further reveal that the two variants rewire EGFR’s internal signal propagation network in distinct ways: L858R diminishes the transmission of structural changes to distal regions of the C-lobe, whereas introducing T790M in addition to L858R partially restores wild-type-like behaviour through compensatory rerouting of early-stage dynamical pathways.

## INTRODUCTION

The epidermal growth factor receptor (EGFR) is a transmembrane protein belonging to the erythroblastic leukaemia viral oncogene B (ErbB) family of tyrosine kinase receptors. Under physiological conditions, EGFR regulates critical cellular processes such as cell proliferation, differentiation and survival.^1^ Upon binding to its natural ligands, such as the epidermal growth factor (EGF), the receptor undergoes dimerisation and autophosphorylation, triggering downstream signalling cascades, including the RAS-RAF-MEK-ERK and PI3K-AKT pathways.^2^ In cancer, particularly in non-small cell lung cancer (NSCLC), mutations in EGFR can lead to its constitutive activation.^3,4,5^ This results in continuous signalling that promotes uncontrolled cell growth, resistance to apoptosis, and tumour progression.^6^ Moreover, activating mutations, such as L858R and T790M, which frequently co-occur in tumours that develop resistance to first- and second-generation tyrosine kinase inhibitors (TKIs) such as gefitinib, represent well-established oncogenic drivers, with the resulting mutant receptors constituting important therapeutic targets for the development of EGFR-targeting drugs.^7^ For example, the L858R/T790M double mutant led to the development of TKI EGFR inhibitors like osimertinib, which caused a marked decline in the prevalence of T790M-positive tumours.^8,9^

EGFR’s structure consists of two modules, an extracellular EGF-binding domain and an intracellular kinase domain, connected by a single-helix transmembrane segment and a juxtamembrane segment (Fig. 1A).^17^ The kinase domain, where most NSCLC-related mutations are located, features a characteristic bilobal architecture, with a smaller N-terminal lobe (N-lobe) composed mainly of β-sheets and a larger C-terminal region (C-lobe) dominated by α-helices (Figs. 1B-1C).^1^ During EGF-induced dimerisation, the two kinase domains come into close proximity, with the C-lobe of one monomer (the *activator*) contacting the N-lobe of the other monomer (the *receiver*). This interaction triggers the conformational changes required to activate the *receiver* monomer.^6^ Once activated, the *receiver* binds and hydrolyses ATP, providing the phosphate groups necessary for the (auto)phosphorylation of multiple tyrosine residues within the tyrosine-rich region located on the intracellular tail (Fig. 1). The phosphorylated residues then act as docking sites for signalling proteins, such as GRB2 and Shc, ultimately modulating downstream pathways that influence gene expression and cellular responses.^18^

**Fig. 1:**
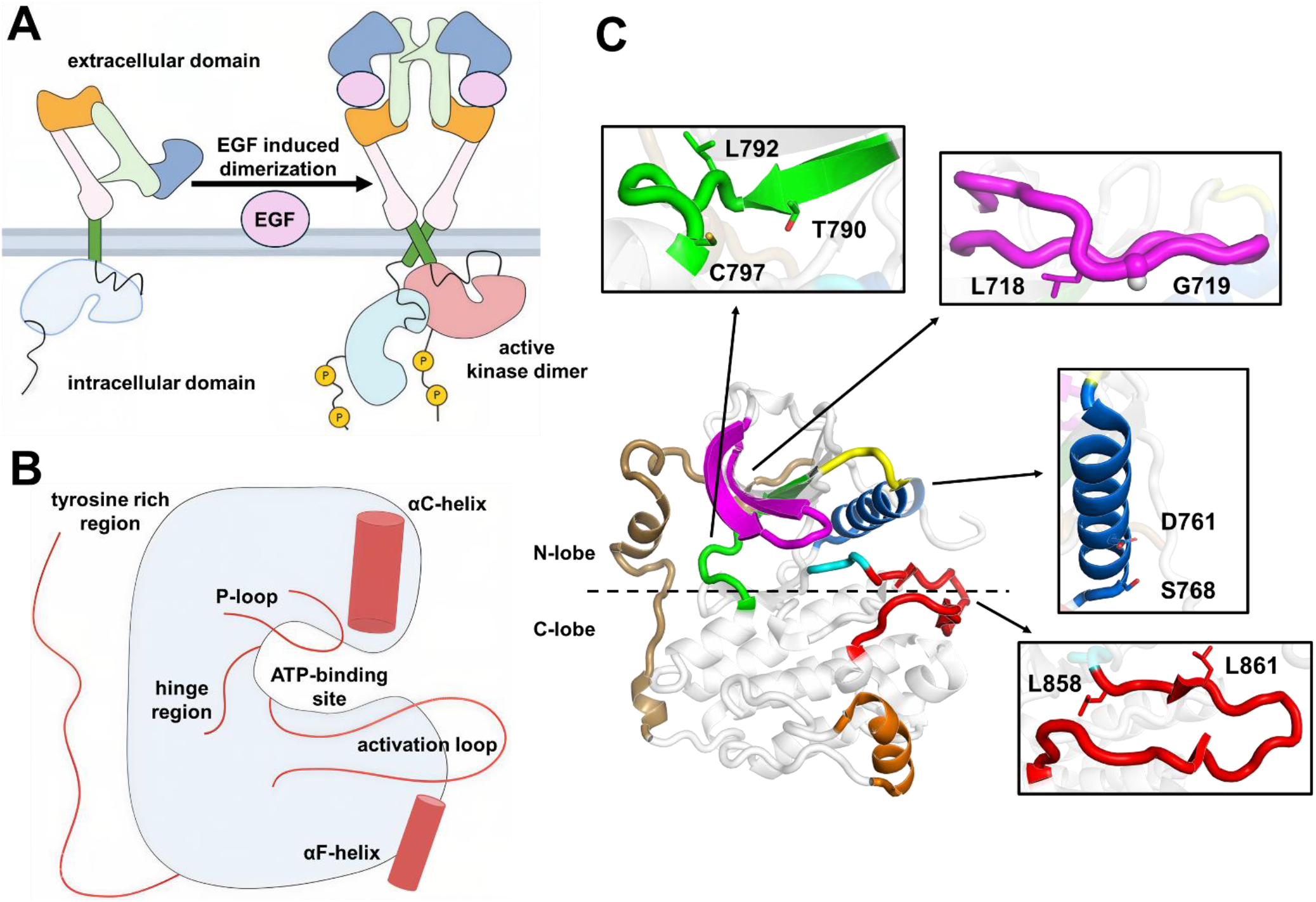
EGFR structure and activation mechanism. A) EGF-induced dimerisation of EGFR. Upon EGF binding, full-length EGFR forms an asymmetric homodimer in which one of the tyrosine kinase subunits (often referred to as the activator; shown in light red in the dimer) allows the other kinase monomer (called the receiver; shown in light blue) to switch from an inactive to an active conformation.^1^ **B) Schematic representation of EGFR’s kinase domain.** The kinase domain consists of N- and C-lobes, with the ATP-binding pocket in the cleft between them. Important structural elements, namely the P-loop, αC-helix, hinge region, activation loop, and αF-helix, are highlighted in red. **C) Three-dimensional structure of the wild-type EGFR kinase domain in the active conformation.** Functional regions and cancer-related mutation sites are mapped onto the crystal structure of wild-type EGFR (PDB ID 8SC7).^10^ Highlighted functional regions include: the P-loop, in pink (residues F712-W731); the αC-helix, in blue (residues S752-D770); the DFG motif (residues 855-857) which belongs to the activation loop, shown in red (residues D855-P877); the hinge region in green (residues L788-C797); the exon-19-encoded segment commonly affected by deletions in cancer shown in yellow (residues T751-K745); the αF-helix in orange (residues P919-G930); and the tyrosine-rich region in brown (980-1019).^11^ Cancer-associated mutation sites (L718, G719,^12,13^ D761, S768,^14^, T790, C797, L858,^15^ L861, and L972^16^) are shown with sticks.

Each kinase monomer contains an ATP-binding site formed by structural elements from both the N- and C-lobes. These include the P-loop, the hinge region, and the highly conserved aspartate-phenylalanine-glycine (DFG) motif in the activation loop (Figs. 1B-1C).^2^ Several distal regions also contribute to the regulation of kinase activity (Figs. 1B-1C), including the αC-helix, the activation loop, and the αF-helix.^14,19^ Experimental evidence indicates that the interface between the *receiver*’s N-lobe and the *activator*’s C-lobe, along with the *receiver*’s P-loop, hinge region, DFG motif, activation loop, and αC-helix, all contribute to the regulation of EGFR’s kinase activity.^20^ The orientations adopted by both the DFG motif and the αC-helix are well established markers of the kinase’s activation state: the αC-helix is displaced outward towards the solvent in the inactive form (Fig. S2B) and in the active state it adopts a conformation closer to the ATP-binding site (Fig. S2A), whereas the DFG motif assumes the typical “DFG-in” position in the active state (Fig. S1A) and a ‘DFG-out’ arrangement in the inactive one (Fig. S1B).^21^ Lastly, a short sequence of residues linking the αC-helix to the P-loop (E746–T751), encoded by exon 19 (Fig. 1C), also contributes to EGFR function, with deletions within this segment leading to some of the most common activating mutations in NSCLC.^22^

Despite extensive experimental and structural data available for EGFR (for recent reviews, see refs 23, 24), including various X-ray and cryo-EM structures of diverse EGFR complexes (e.g. PDB IDs 9QXN,^25^ 9S3X,^26^ 9GL9,^27^ and 9H42^26^), the mechanisms by which functional and regulatory sites communicate remain poorly understood. To address this, numerous computational studies have explored EGFR’s functional dynamics, using methods ranging from molecular mechanics-molecular dynamics (MM-MD) simulations to metadynamics and hybrid quantum mechanics/molecular mechanics (QM/MM) approaches. For example, using extensive equilibrium MD simulations, Arkhipov *et al.* showed that basic residues in the N-lobe of the *receiver* play a crucial role in stabilising both the active and inactive receptor conformations by interacting with the membrane.^20^ Dixit *et al.* used equilibrium MD simulations combined with essential dynamics and principal component analysis to investigate long-range connections in EGFR, showing that communication between the αC-helix, the activation loop, and the αF-helix is tightly linked to the activity of the kinase domain.^19^ Metadynamics simulations by Sutto *et al.* pointed to the crucial role of the activation loop and αC-helix in regulating EGFR activity, showing that coordinated motions of these regions favour active conformation in variants harbouring the cancer-associated L858R and T790M mutations, whereas the wild-type protein preferentially adopts an inactive conformation.^28^ Kannan *et al.* combined extensive equilibrium MD simulations with binding assays to investigate how mutations influence the formation of a potentially druggable allosteric pocket, showing that variants such as L858R, L858R/T790M, and L861Q induce structural changes that destabilise the inactive state of EGFR and promote a transient intermediate in which the allosteric pocket becomes fully formed and accessible to ligand binding.^29^ Finally, hybrid QM/MM simulations by Callegari *et al.* investigated the impact of the P-loop L718Q mutation, which is linked with resistance to third-generation covalent TKIs; surprisingly, their results suggested that, rather than changing ligand affinity or the energetics of covalent bond formation, L718Q stabilises the inhibitor in a non-reactive binding pose. This non-reactive orientation prevents the electrophilic warhead of third-generation drugs from aligning with C797, thereby blocking the covalent bond formation required for irreversible inhibition and ultimately permitting the kinase to remain catalytically active.^13^

Despite tremendous progress in recent years in elucidating EGFR’s structure, dynamics, and function, mechanistic understanding of how conformational signals propagate within the receptor remains incomplete. In particular, the internal pathways that mediate long-range communication within the kinase domain, and how cancer-associated mutations at distal positions (i.e., located far from the ATP binding site) modulate EGFR’s conformational landscape and activity, are still not fully understood. Better understanding of the functional mechanisms and effects of distal mutations may aid in developing new (e.g. allosteric) EGFR therapeutics.

Here, we apply the emerging dynamical-nonequilibrium MD (D-NEMD) approach^30,31^ to elucidate the dynamic mechanisms underlying allosteric modulation in EGFR and to dissect how cancer-associated mutations perturb the kinase’s functional dynamics. D-NEMD, which combines equilibrium and nonequilibrium simulations,^32^ has shown great potential to address a variety of biomolecular questions, including tracking the transmission of structural changes,^33^ identifying allosteric effects,^34^ predicting sites that modulate enzyme activity,^35^ and evaluating the impact of pH changes^36,37^ in proteins. In this work, D-NEMD, exploiting instantaneous ligand removal from the nucleotide binding site as the external perturbation, is used to probe how structural changes propagate within EGFR’s wild-type kinase monomer, and how two of the most prevalent NSCLC mutations, L858R and L858R combined with T790M, affect the protein’s dynamic responses and internal communication pathways.^15^ L858R, located in the activation loop of the kinase domain (Fig. 1C), enhances its *k*_cat_ by more than 50 times with respect to the wild-type and stabilise the kinase’s active conformation,^28,38^ although it reduces ATP affinity by approximately 30 times.^38^ T790M, within the ATP-binding pocket (Fig.1C), by itself mainly increases catalytic efficiency by increasing *k*_cat_, with only a small change in ATP affinity, thereby conferring strong resistance to ATP-competitive inhibitors.^38,39^ Notably, the combined effect of L858R and T790M is synergistic, with the double mutant generally displaying a similar ATP affinity as the wild type but an ∼18-fold increase in *k*_cat_.^38,39^ Therefore, these differences in catalytic properties and ATP affinity suggest that the wild-type and L858R systems may exhibit distinct internal communication pathways and conformational dynamics, whereas L858R/T790M double mutant may retain dynamical features characteristic of both systems.

Our D-NEMD results reveal distinct allosteric networks and dynamic behaviours between the wild-type and mutants, indicating differences in the regulation of functional regions and subtle variations in their internal communication networks. In the wild type, these networks involve regulatory structural elements, including the P-loop, hinge region, activation loop, αC-helix, an exon-19 segment, the tyrosine-rich region, and the N-lobe interface that engages the C-lobe of the *activator* monomer upon dimerisation. The L858R and L858R+T790M mutations reconfigure these communication pathways, with the largest differences relative to the wild type located in the N-lobe, αC-helix, activation loop, and αF-helix. The findings show how oncogenic substitutions reshape signal propagation from the ATP site and alter EGFR’s regulatory architecture.

## RESULTS AND DISCUSSION

Equilibrium MD and D-NEMD simulations were used to probe the functional dynamics of the wild-type EGFR kinase domain and to assess how two of the most prevalent cancer-associated activating mutations, L858R and T790M,^15,40^ affect the domain’s behaviour.

To characterise how these mutations influence kinase dynamics, extensive long-timescale equilibrium MD simulations were performed, followed by a large D-NEMD simulation set (comprising > 1300 nonequilibrium trajectories), yielding a total of 19 μs of simulation. The starting structures for the equilibrium simulations were derived from the crystal structure of the monomeric EGFR kinase domain in its active conformation, with the non-covalent BLU-945 inhibitor (also known as tigozertinib; PDB ID 8D76)^41^ bound at the ATP site. This structure contains the L858R and T790M substitutions, as well as the crystallisation-facilitating V948R mutation.^42^ V948R has been shown experimentally not to affect substrate affinity or enzymatic activity, indicating that it does not alter the biochemical properties of the monomeric EGFR kinase domain.^42^ We therefore retained V948R in all models to preserve (as much as possible) the experimentally resolved structural environment of the crystal template.

Using this crystal structure, we constructed three EGFR kinase-BLU-945 complexes (details in Materials and Methods): (*i*) a wild-type-like system, in which L858R and T790M were reverted to their original wild-type sequences, while retaining V948R (hereafter referred to as *wild-type*); (*ii*) a mutant system containing L858R and V948R, with residue 790 restored to threonine (labelled *L858R*); and (*iii*) a mutant system harbouring the L858R, T790M and V948R substitutions (labelled *L858R+T790M)*. Experimental evidence indicates that the BLU-945 inhibitor preferentially binds to mutant EGFR variants.^41,43^

For each EGFR-inhibitor complex, five 500-ns equilibrium MD simulations were performed. All systems remained stable throughout the trajectories and reached equilibrium after ∼50 ns (Fig. S5). Cα root-mean-square deviations (RMSD) and principal component analysis (PCA) were used to evaluate equilibration and sampling across replicates for wild-type and mutant systems. PCA also shows overlap in the principal motions across all three systems: wild-type and mutant complexes exhibit similar dominant collective movements (Fig. S7). The first two principal components, capturing the largest-amplitude and most correlated motions across the systems, were primarily associated with P-loop movements, αC-helix bending, and unfolding of the tyrosine-rich region (Figs. S4 and S7). The BLU-945 inhibitor remained stably bound to the ATP site in both wild-type and mutant systems (Fig. S10).

Root-mean-square fluctuations (RMSF) were calculated for each Cα atom (Fig. S6) to assess local flexibility across systems and identify differences in their dynamics. Overall, RMSF profiles were similar among replicates of the same system, with only variations observed in the αC-helix and in the solvent-exposed tyrosine-rich region (residues V980D1014). Between systems, the largest RMSF differences were located in the αC-helix, which is more mobile in the *wild-type* and *L858R+T790M* variant, but more rigid in the *L858R* mutant.

Statistical correlations between position 858 and all the other Cα atoms for the *wild-type* and mutants were determined to identify regions of the protein whose motions are coupled to the mutation site and how these couplings are affected by the mutations. This analysis shows that, in the equilibrium simulations, the mutations do not drastically change the overall coupling between the motions of residue 858 and the remaining residues (Fig. S11).

As mentioned above, the DFG motif and αC-helix orientations are well established markers of kinase activation.^21,28^ The DFG conformation was monitored by measuring the distance between the phenyl ring of F856 and the Cα of M766 (Fig. S8), a metric that distinguishes the active DFG-in from the inactive DFG-out configuration. As shown in Fig. S8, all simulated systems maintained the DFG-in conformation throughout the equilibrium simulations, indicating that the complexes retain structural features characteristic of active EGFR. αC-helix behaviour was studied by measuring the distance between the salt bridge formed by K745 and E762. As shown in Fig. S9, the average distance resembles a semi-active conformation already described by Sutto *et al.*, ^28^ where the αC-helix resembles an inactive-like EGFR conformation, as the distance between K745 and E762 is not suitable for a salt bridge, but is still much lower than the distance observed in inactive EGFR.

To verify that the crystallisation-facilitating V948R substitution does not alter the intrinsic dynamics of the kinase domain, we simulated a fourth system (hereafter labelled as *control*), in which all three mutations (L858R, T790M, and V948R) were reverted to the original sequence. Comparison of the behaviour of the *wild-type* and *control* systems in both equilibrium and nonequilibrium simulations confirmed that V948R has a negligible effect on kinase-domain dynamics (Figs. S4-S9 and Figs. S14-S19).

### Dynamical behaviour of *wild-type*

D-NEMD probes dynamical responses from differences between equilibrium and nonequilibrium trajectories.^30,32^ From the equilibrium MD simulations above, 450 short (5 ns) nonequilibrium MD trajectories were generated for the *wild-type, L858R, L858R+T790M*, and *control* systems (i.e., 1800 total) to characterise the kinase’s dynamical behaviour and map the internal networks that coordinate its functional motions. For this, an ensemble of 450 conformations per system (90 per replicate) was extracted from the equilibrium trajectories and used as starting points for the nonequilibrium MD simulations, in which BLU-945 was instantaneously removed from the ATP binding pocket (Figs. S3 and S12). These simulations capture the evolution of the kinase’s structural and dynamic response as it adapts to an empty ATP site. BLU-945 removal served as a (well-defined and controlled) external perturbation that pushed the system out of equilibrium, creating the driving force required to expose the internal communication pathways engaged as the protein gradually relaxed back toward a (new) equilibrium state. BLU-945 removal triggered an initial structural response in the ATP pocket that, over time, propagated through the protein as a coordinated sequence of conformational changes.

In addition to the nonequilibrium trajectories, 450 “null perturbation” simulations were conducted for the *wild-type*, *L858R*, *L858R+T790M,* and *control* systems (see “Materials and Methods” in Supporting Information; Fig. S12) to distinguish intrinsic equilibrium fluctuations from perturbation-driven responses. The “null perturbation” analysis^44^ entails the subtraction of equilibrium motions from D-NEMD responses, thereby reducing noise and distinguishing dynamical signals from intrinsic equilibrium movements of the system.^34,35,36^

Dynamical responses were calculated using the Kubo-Onsager relation (see “Materials and Methods” in Supporting Information for a detailed description; Fig. S12). The amplitude and directionality of the Cα responses were obtained by determining the difference in atomic positions between paired equilibrium and nonequilibrium trajectories at equivalent time points, and subsequently averaging across all 450 replicates per system. The resulting signals were then refined using the “null perturbation” controls to remove equilibrium fluctuations and isolate the perturbation-induced changes (Figs. S12 and S15-S18), following the approach described previously.^31,36^ The standard error of the mean (SEM) was used to assess convergence of the Cα responses, ensuring that the analysis reflected meaningful dynamical changes (Fig. S12).

Analysis of the evolution of *wild-type* conformational rearrangements following BLU-945 removal reveals the complex mechanical coupling between different structural elements involved in signal transmission from the ATP site to distal regions (Figs. 2 and S15). As shown in Fig. 2, the cascade of structural changes initiated by ligand removal propagates outward gradually from the ATP pocket, ultimately affecting functionally relevant regions far from the perturbation site. These include the N-lobe, the αC-helix, the αF-helix, the activation loop, and the distal tyrosine-rich region.

**Fig. 2:**
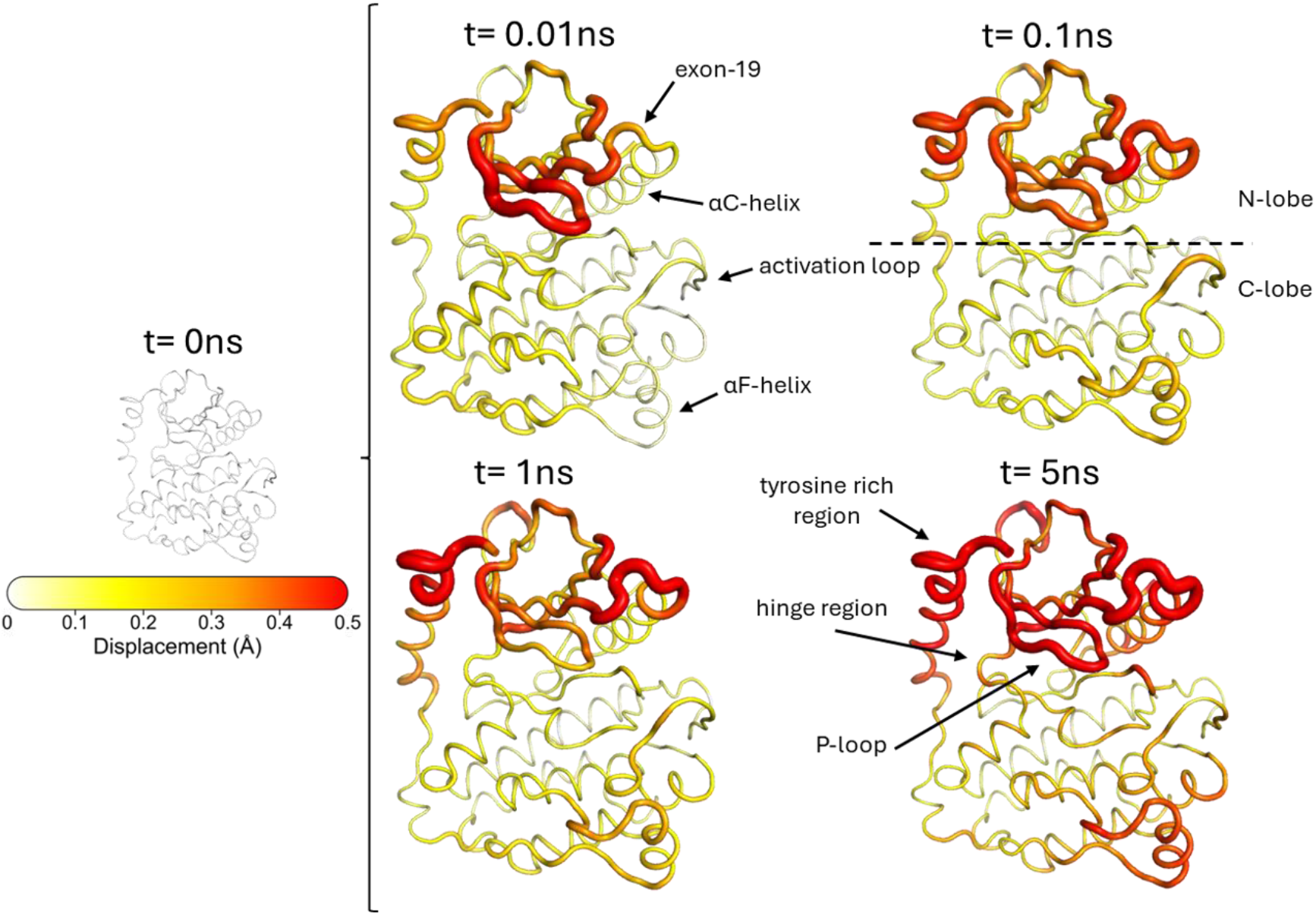
Dynamical response of *wild-type* kinase to inhibitor removal. The average Cα displacements at 0, 0.01, 0.1, 1, and 5 ns following BLU-945 removal are mapped onto the *wild-type* average structure from the equilibrated part (from 50 to 500 ns) of the equilibrium MD simulations. The cartoon thickness and structure colours (scale on the left) depict the magnitude of the protein’s average response to ligand removal.

As expected, the earliest structural changes arise in motifs closest to the ATP site, including the P-loop and hinge region, which undergo rapid rearrangement immediately after BLU-945 removal. Clear responses are observed as early as 0.01 ns, particularly within the P-loop and exon-19 region. From these sites, the signal progresses toward the tyrosine-rich region, the activation loop, and the hinge region, ultimately reaching the αF-helix in the C-lobe. The N-lobe, which includes important functional regions such as the P-loop and αC-helix, shows greater sensitivity to ligand removal than the C-lobe, displaying more pronounced displacements (Fig. 2).

Many of the most common cancer-associated mutation sites (including positions 746–750, 790, 797 and 858) fall within, or in direct contact with, the regions that D-NEMD analysis identifies as exhibiting the most pronounced responses to BLU-945 removal (Figs. 3A and S16). These positions include T790, C797, and L792 in the hinge region; L861 and L858 in the activation loop; G735 in the N-lobe; D761 and S768 in the αC-helix; and L718 and G719 in the P-loop, which map onto the communication pathways through which structural signals propagate from the ATP site (Fig. 3A). This overlap between the D-NEMD identified communication routes and clinically relevant mutation hotspots is exemplified by exon-19 deletions (typically involving residues 746– 750) and the L858R substitution in the activation loop, which together account for ∼ 85% of all activating EGFR mutations in NSCLC.^15,16^ Exon-19 mutations promote ligand-independent kinase activation and uncontrolled cell proliferation by enabling tighter packing of the αC-helix on the ATP site, thereby facilitating autophosphorylation.^45,46^ These mutations occur on highly responsive segments of the pathways that relay structural changes from the ATP pocket; thus, they are expected to have substantial functional consequences, consistent with experimental observations.

**Fig. 3:**
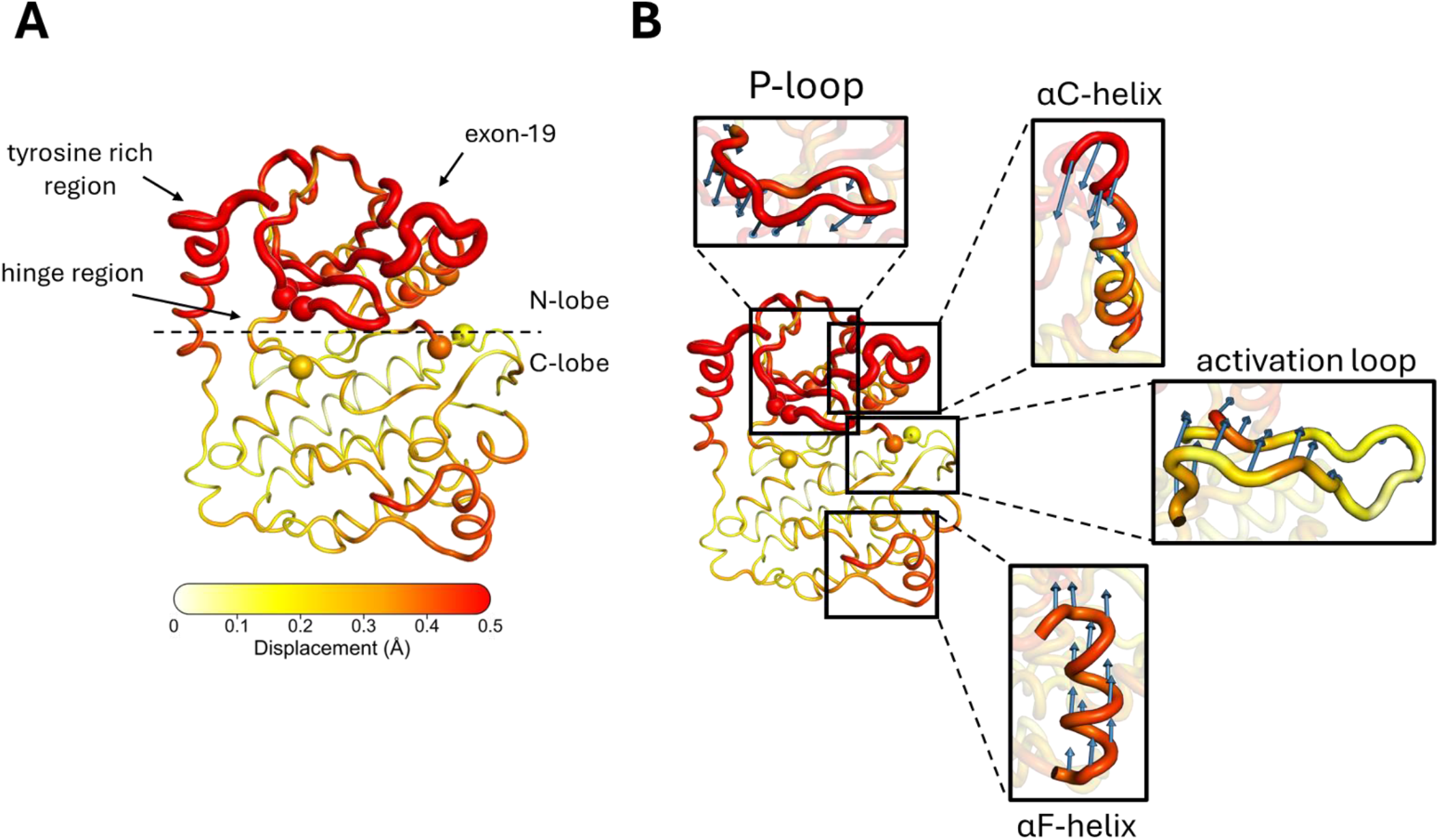
Mutation sites identified by D-NEMD and direction of the structural response of *wild-type* to BLU-945 removal. **A)** The average Cα displacements are mapped onto the *wild-type* average structure, with some of the most relevant functional regions labelled. The average displacement magnitudes are represented on a white-yellow-orange-red scale. The positions of residues linked to cancer-related mutations, including two of the most common NSCLC substitutions, L858R and T790M, are highlighted as spheres. The residues highlighted are T790, C797, and L792 in the hinge region; L861 and L858 in the activation loop; G735 within the N-lobe; D761 and S768 in the αC-helix; and L718 and G719 in the P-loop **B)** Cα displacement vectors after 5 ns mapped onto *wild-type* average structure, with detailed view of the P-loop, αC-helix, activation loop, and αF-helix regions. Vectors with lengths ≥ 0.2 Å are displayed as blue arrows (scaled by a factor of 5).

The T790M “gatekeeper” substitution is positioned within the pathways identified by D-NEMD. This mutation is a known driver of resistance to first- and second-generation TKIs, such as erlotinib and gefitinib,^38,47^ and has been detected in 50–60% of NSCLC patients who develop resistance to TKIs after an initial positive therapeutic response.^48^ Its clinical prevalence prompted the development of third-generation TKIs, such as osimertinib, which were specifically engineered to overcome T790M-mediated resistance by selectively and irreversibly targeting this mutant kinase while sparing the wild-type EGFR.^49,50^ The strategic positioning of T790M within the hinge region, a key node along the D-NEMD communication pathways that transmit structural changes, likely contributes to its profound functional consequences, especially in combination with other cancer-associated mutations such as L858R.^38^

D-NEMD analysis also shows the direction of the motions involved in the response process (Fig. 3B). In the *wild-type* system, five nanoseconds after BLU-945 removal, residues F712-W731 within the P-loop (positioned above the ATP site) exhibit a clear inward movement, consistent with the closing of the ATP pocket (Fig. 3B). Concurrently, the upper part of the αC-helix (S752-D770) displays a similarly well-defined motion that follows the P-loop rearrangement, displaying an outward rotation (Fig. 3B) indicative of the early transition from an active to an inactive-like conformation, as observed in the K745-E762 distance analysis during the equilibrium simulations (Fig. S9). Finally, the activation loop (D855-P877) and the αF-helix (P919-G930) both shift upward, indicating coordinated long-range responses in regions distal to the ATP site (Fig. 3B).

### Effects of L858R and L858R+T790M mutations

D-NEMD was next applied to examine if, and if so, how two of the most prevalent activating mutations in NSCLC, L858R and L858R+T790M, reshape the kinase’s dynamical behaviour and its internal communication networks (Figs. 4-5 and S17-18). For this, following the same protocol used for the *wild-type* system, 450 short (5 ns) nonequilibrium simulations and 450 short (5 ns) “null perturbation” simulations were generated for each mutant by instantaneously removing BLU-945 from the ATP pocket.

**Fig. 4:**
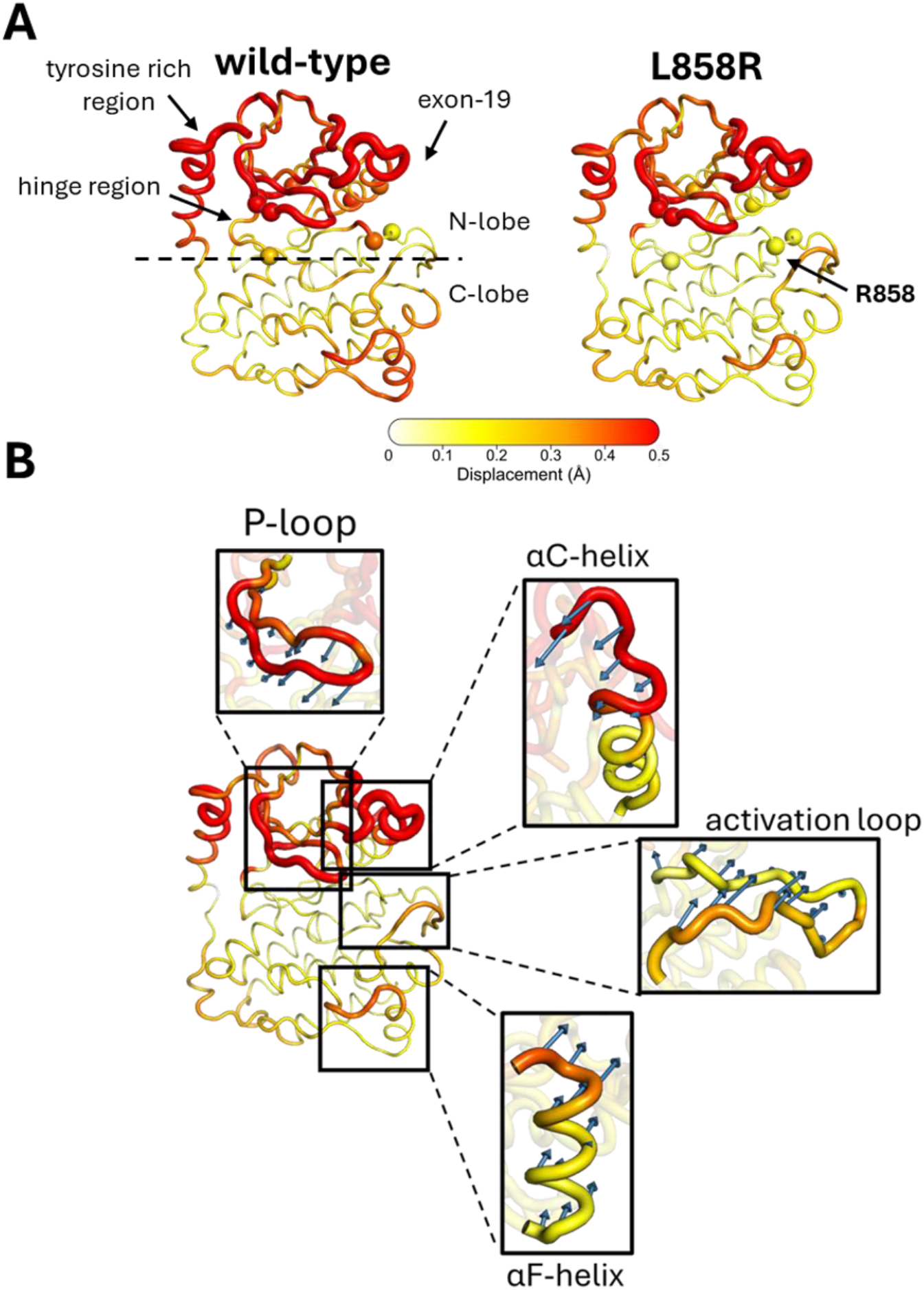
Mutation sites identified by D-NEMD and dynamical response of *L858R*. **(A) Average displacements for *wild-type* (left panel) and *L858R* (right panel) at 5 ns following BLU-945 removal.** The average Cα displacements are mapped onto the *wild-type* average structure, calculated from the equilibrium simulations. In this figure, the cartoon thickness and structure colours (scale on the bottom of Fig. 4A) indicate the amplitude of the proteins’ response to BLU-945 removal. Residues linked to cancer-related mutations, including two of the most common NSCLC substitutions, L858R and T790M, are highlighted as spheres. These include T790, C797, and L792 in the hinge region; L861 and L858 in the activation loop; G735 within the N-lobe; D761 and S768 in the αC-helix; and L718 and G719 in the P-loop. **(B) Direction of the structural response of *L858R* at 5 ns following BLU-945 removal.** Cα displacement vectors after 5ns mapped onto the *wild-type* average structure, with a detailed view of the P-loop, αC-helix, activation loop, and αF-helix regions. The average Cα displacement vectors at t = 5 ns are shown as blue arrows. Vectors with lengths ≥ 0.2 Å are displayed as arrows (scaled by a factor of 5) and mapped onto the *wild-type* average structure. The average displacement magnitudes are represented on a white-yellow-orange-red scale.

**Fig. 5:**
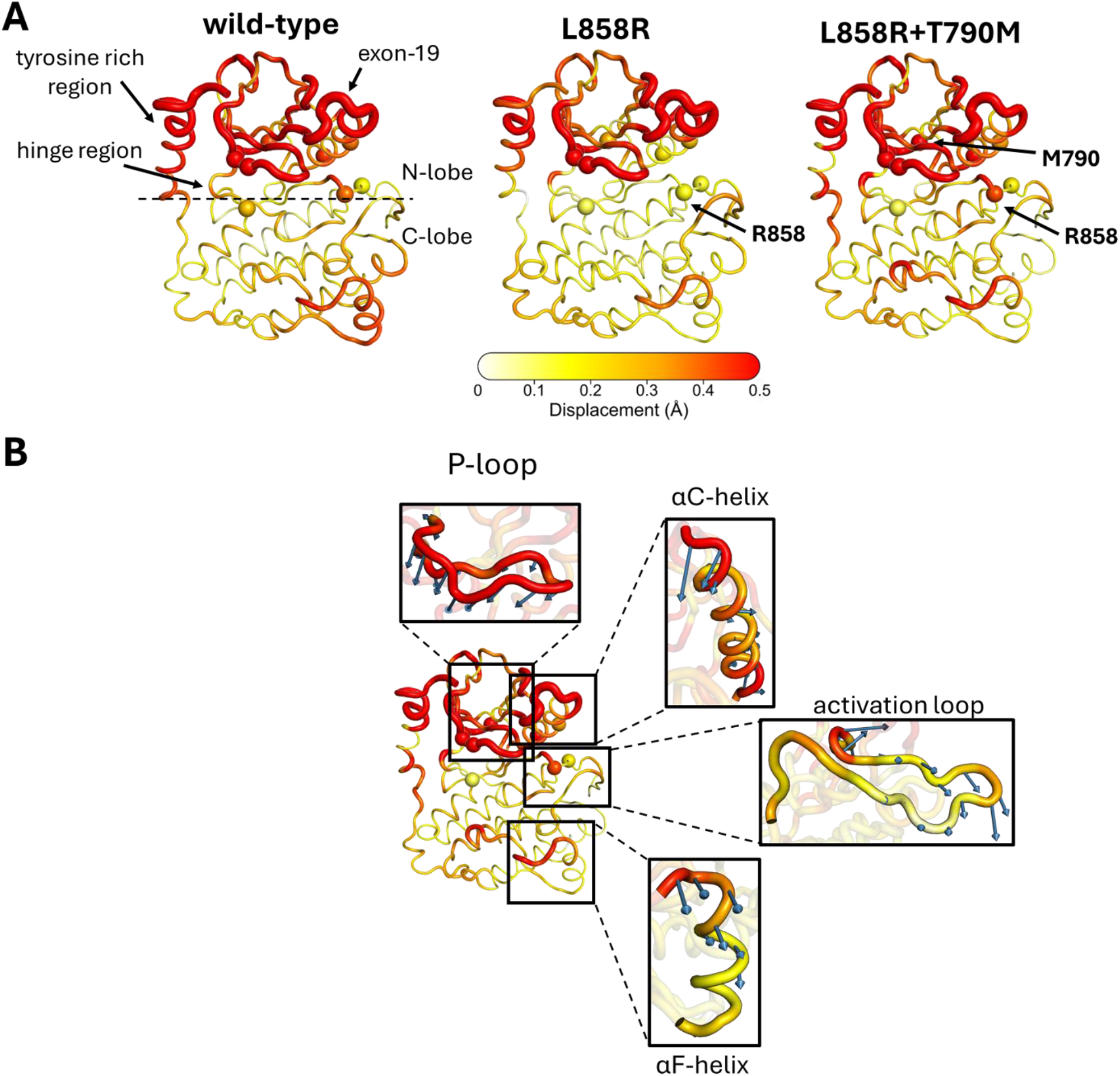
Mutation sites identified by D-NEMD and dynamical response of *L858R+T790M*. **(A) Average displacements of wild-type (left panel), L858R (middle panel), and *L858R+T790M* (right panel) at 5 ns following BLU-945 removal.** The average Cα displacements are mapped onto *wild-type* average structure. In this figure, the cartoon thickness and structure colours (scale on the bottom) indicate the amplitude of the proteins’ response to BLU-945 removal. Residues linked to cancer-related mutations, including two of the most common NSCLC substitutions, L858R and T790M, are highlighted as spheres. These include T790, C797, and L792 in the hinge region; L861 and L858 in the activation loop; G735 within the N-lobe; D761 and S768 in the αC-helix; and L718 and G719 in the P-loop. **(B) Direction of the structural response of *L858R+T790M* at 5 ns following BLU-945 removal.** Displacement vectors after 5ns mapped onto the *wild-type* average structure, with a detailed view of the P-loop, αC-helix, activation loop, and αF-helix regions. Vectors with lengths ≥ 0.2 Å are displayed as arrows (scaled by a factor of 5) and mapped onto the *wild-type* average structure. The average displacement magnitudes are represented on a white-yellow-orange-red scale.

As in the *wild-type*, BLU-945 removal in the *L858R* mutant initiates a cascade of structural changes that propagates through the kinase domain, initially impacting regions near the ATP site, including the P-loop, the αC-helix, and after 5 ns, expanding across the rest of the N-lobe, reaching the tyrosine-rich region (Fig. 4A). However, in *L858R*, the C-lobe, including the activation loop and αF-helix, display markedly attenuated responses relative to the *wild-type* following inhibitor removal (Fig. 4A).

A further striking difference between *L858R* and *wild-type* lies in the behaviour of the αC-helix, the tyrosine-rich region, and the hinge region, all of which show noticeably reduced structural responses in the mutant. In the *wild-type,* these elements are among the earliest and most responsive segments of the internal network connecting to the ATP site; however, in *L858R,* each of these regions shows substantially reduced sensitivity to the ATP-site perturbation, suggesting that the initial conformational signal fails to propagate as efficiently along these pathways (Fig. 4A).

The directionality of the structural response in *L858R* was also examined and compared with the *wild-type* (Fig. 4B). Although the overall magnitude of the *L858R* response after 5 ns is generally reduced relative to the *wild-type*, particularly within the C-lobe, the direction of motion in key early-responding regions, such as the P-loop and exon-19 segment, remains largely similar to the *wild-type* behaviour (Figs. 3 and S15).

These differences suggest that *L858R* compromises the coordinated flow of structural changes across the C-lobe, weakening its functional connection to the ATP site. The altered dynamical behaviour of the αC-helix is particularly striking (Fig. 4B), with its diminished responsiveness likely disrupting long-range signal transmission across the receptor, leading to a breakdown of the coordinated, domain-wide connections present in the *wild-type*.

Overall, our D-NEMD findings highlight how *L858R* rewires the connections between the ATP cleft and distal functional and regulatory regions, thereby influencing active-inactive transitions and likely weakening long-range inter-domain communication (e.g. between the C-lobe of one kinase monomer and the N-lobe of the other upon dimerisation). Notably, the fact that D-NEMD, despite using fully unbiased simulations, captures these mutation-dependent changes demonstrates the method’s ability to detect functionally meaningful dynamical alterations and provide new mechanistic insight into how proteins respond to external triggers.

D-NEMD was also used to examine the combined effect of L858R and T790M, using the same protocol described above for the *wild-type* and *L858R* systems (Figs. 5 and S19). Experimental data indicate that the ∼30-fold reduction in ATP affinity caused by the single L858R mutation is reversed by the introduction of T790M, with the double mutant generally showing an affinity comparable to that of the wild-type kinase.^38^ Kinetic measurements further indicate that although L858R can increase catalytic turnover by up to ∼50-fold relative to wild type, this enhancement is reduced to ∼18-fold in the double mutant.^38,39^ Together, the T790M and L858R mutations produce a hyperactive kinase that combines features from both individual mutants, retaining similar ATP affinity as the wild type while maintaining increased activity.^38,39^ Consistent with this, the D-NEMD responses of *L858R+T790M* to BLU-945 removal exhibit a hybrid behaviour, combining the patterns of response of both *wild-type* and *L858R* (Fig. 5A). For example, the activation loop, hinge region, and tyrosine-rich segment show structural changes closely resembling those of the *wild-type*, whereas the αF-helix behaves more like *L858R*. Additionally, the αC-helix and the C-lobe show an intermediate level of sensitivity to BLU-945 removal, with their response amplitudes falling between the strongly responsive *wild-type* and the muted behaviour observed in *L858R* (Fig. 5A) This observed profile further reinforces the view that *L858R+T790M* blends dynamical traits from both parent systems.

In *L858R+T790M,* as in the *wild-type* (and also *L858R)*, BLU-945 removal initiates a cascade of conformational changes that originate in the ATP site and subsequently spread to nearby elements, including the P-loop and the αC-helix, before reaching more distal regions such as the activation loop and the αF-helix (Fig. S18). However, there are differences in the timings of the earliest events between systems: for example, in *L858R+T790M*, the αC-helix is the first element to react within the initial 100 ps following BLU-945 removal, followed by a pronounced displacement of the P-loop at ∼1 ns; in contrast, in *wild-type*, the P-loop exhibits the earliest response to inhibitor removal, with αC-helix rearrangements occurring later (Figs. S15-S19). Despite these early differences in the order of the events, by 5 ns, the *wild-type* and *L858R+T790M* display comparable P-loop and αC-helix displacements, indicating convergence toward a similar dynamical response in these motifs. This means that although the initial pathways of structural change propagation differ, *L858R+T790M* ultimately adopts *wild-type*– like conformational behaviour in these structural elements. The altered early-stage response observed in *L858R+T790M* may reflect a compensatory allosteric mechanism that restores aspects of wild-type behaviour, thereby reconciling its L858R-driven hyperactivity with the recovered ATP affinity characteristic of the *wild-type*. ^38^

Another important distinction between *L858R+T790M*, *wild-type,* and *L858R* responses lies in the directionality of their motions (Fig. 5B). For example, although the activation loop and αF-helix display broadly similar displacement magnitudes across the three systems after 5 ns, the orientation of these movements differs markedly, undergoing a predominantly downward motion toward the solvent in *L858R+T790M,* whereas shifting upwards toward the activation loop in the *wild-type* and *L858R* systems.

Analysis of the hydrogen-bonding network formed by R858 with residues in the activation loop (D855-P877) and αC-helix (S752-D770) during both equilibrium and nonequilibrium simulations (Figs. S20-21) showed a marked difference between the *L858R* and *L858R+T790M* systems, with the latter forming interactions with more residues across both key regions. This finding suggests that position 858 acts as an important communication node linking the N- and C-lobes, providing a possible mechanistic explanation for the different behaviour of *L858R*.

Overall, while the displacement amplitudes of several elements in *L858R+T790M* eventually converge toward either *wild-type*-like or *L858R*-like values, their directional vectors often do not, instead retaining unique directional signatures, highlighting that subtle differences in early response pathways evolve into distinct longer-timescale dynamical patterns.

## CONCLUSION

In this study, we investigate the intricate allosteric network within the kinase domain of the EGFR to map functionally relevant communication networks and understand how two of the most common, clinically relevant EGFR activating mutations reshape these internal pathways. By systematically comparing the responses of *wild-type*, *L858R*, and *L858R+T790M* variants to the instantaneous removal of the BLU-945 inhibitor from the ATP site, we provide a mechanistic description of mutation-dependent allosteric regulation and its impact on the kinase functional dynamics. We further show that the allosteric networks mapped in the *wild-type* are strongly associated with sites of oncogenetic variation, with 83% of the mutations observed in NSCLC patients (Fig. S16) falling either directly within these networks or within second-shell contact distance of the mutated residues (Fig. S16).

Our D-NEMD analyses revealed that, in the *wild-type* system, inhibitor removal initiated a rapid and highly coordinated cascade of structural changes involving the P-loop, hinge region, exon-19 segment, αC-helix, activation loop, αF-helix, and the tyrosine-rich region. Together, these elements form an interconnected dynamic network that links the ATP-binding cleft to distal regions of the protein, with the N-lobe emerging as a central hub for signal propagation across all studied systems. Additionally, comparison of the dynamical responses of the *wild-type* and mutants revealed a common network for allosteric communication through which local perturbations are propagated. Disease-associated mutations do not completely rewire this network; rather, they reshape (to different extents) the timing, magnitude, and directionality of structural signal transmission within it.

In the *wild-type*, BLU-945 removal triggered a well-coordinated sequence of structural rearrangements that collectively modulated the accessibility, geometry, and dynamics of the ATP site. Following perturbation, the P-loop moved inward while the αC-helix shifted outward (in a motion that was consistent with the onset of an active-to-inactive conformation transition), whereas the activation loop and αF-helix moved upward toward the N-lobe, forming a concerted response that efficiently propagated the structural changes through the protein, reflecting the integrity of the *wild-type* communication network. Notably, several regions showing the strongest responses to ligand removal aligned with major oncogenic and cancer resistance-associated mutation positions, including sites in exon-19, T790, C797, and L858, underscoring their central role in regulating the kinase’s functional dynamics.

D-NEMD also highlighted how the L858R mutation redefined the kinase’s functional behaviour: although the N-lobe (particularly the P-loop) retained broadly similar response patterns to those of the *wild-type*, the C-lobe became significantly more stable, resulting in altered communication routes and changes in response directionality. The mutation dampened the transmission of structural changes from the ATP site to distal elements, including the αF-helix, tyrosine-rich region, and αC-helix, several of which are located at, or contribute to, the kinase dimer interface (Fig. 6). The reduced responsiveness of these interface-forming elements likely restricts signal propagation between monomers, suggesting a mechanistic basis for how the mutation disrupts long-range allosteric communication within EGFR.

**Fig. 6:**
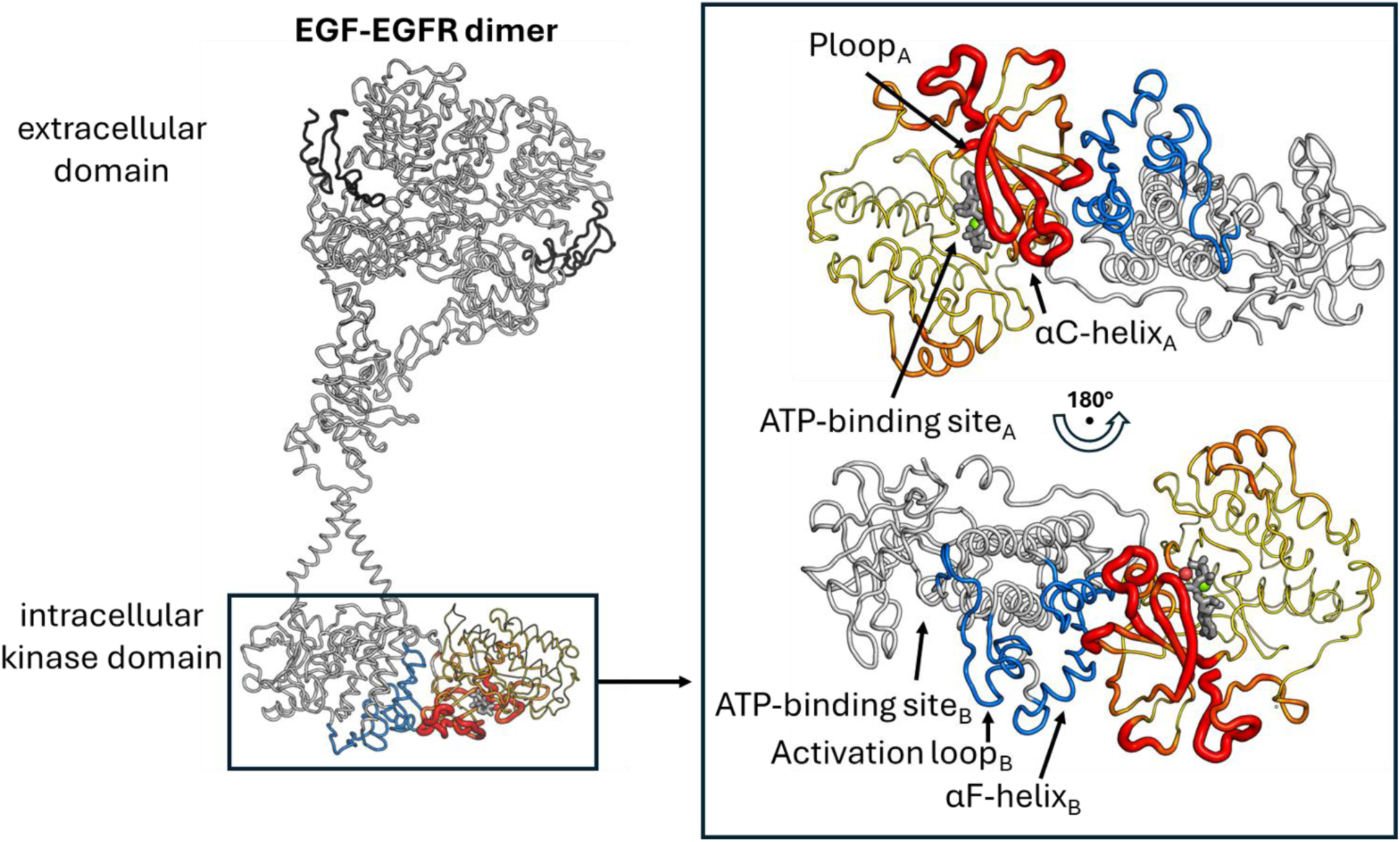
Structural response of the *wild-type receiver* kinase domain to inhibitor removal mapped onto the full-length EGF-EGFR dimer. The average Cα displacements measured 5 ns after BLU-945 removal from the ATP-binding site of *wild-type* EGFR (Fig. S15) are mapped onto the *receiver* kinase (monomer A) of the full-length EGF-EGFR model developed by Arkhipov *et al*.^20^ The cartoon thickness and structure colours of the *receiver* kinase reflect its average Cα displacements according to the scale shown at the bottom of the figure (see the Fig. 2 caption for a detailed description of how the structural responses of the monomer were calculated). Each EGF and EGFR monomer is shown in dark and light grey, respectively. Regions of the *activator* kinase (monomer B), including the αF-helix and activation loop segment, that directly interface with the *receiver*’s domain elements exhibiting strong responses to BLU-945 removal are highlighted in blue. The ATP molecule bound to the *receiver*‘s active site is shown as grey sticks, and its coordinated Mg²^+^ ion and water oxygen are depicted as green and red spheres, respectively.

Remarkably, *L858R+T790M* displayed a hybrid dynamical profile, combining response features of both *wild-type* and *L858R* responses. This behaviour aligns well with experimental evidence showing that the double mutant produces a hyperactive kinase that retains wild-type-like ATP affinity while preserving the elevated catalytic activity introduced by L858R. Consistent with this mixed biochemical phenotype, the P-loop, activation loop, hinge region, and the tyrosine-rich segment in *L858R+T790M* responded to BLU-945 removal in a manner closely resembling the *wild-type*. In contrast, the αF-helix exhibited a response similar to *L858R*. Additionally, although several regions of *L858R+T790M* largely recovered *wild-type*-like dynamical behaviour on the nanosecond timescale, the initial response to inhibitor removal proceeded through a distinct propagation pathway. The altered ordering and directionality of these early events suggest that T790M does not simply reverse the effects of L858R; instead, it reconfigures the kinase’s internal communication landscape, enabling alternative routes for structural signal transmission that restore more effective long-range coupling. Such compensatory rewiring provides a mechanistic explanation for the experimentally observed recovery of ATP affinity and enhanced catalytic efficiency in this double mutant.

More broadly, this work highlights the remarkable plasticity and robustness of the EGFR allosteric communication network, showing how mutations can reorganise internal signal-transmission pathways to preserve, restore, and even enhance functional output while simultaneously introducing new layers of allosteric coupling within the kinase domain. In the context of the full-length receptor dimer, such changes are likely to extend beyond the kinase domain itself. By altering the propagation of structural changes to elements located at the dimer interface, oncogenic mutations may influence inter-domain communication and the transmission of information across the receptor, ultimately influencing distal regions of EGFR and potentially contributing to altered allosteric regulation or activation dynamics (Fig. 6).

Finally, this work establishes D-NEMD as a versatile tool for mapping allosteric pathways in kinases and dissecting the dynamical consequences of cancer-related mutations, an essential step toward understanding how structural perturbations translate into altered cellular behaviour and for informing the design of more effective therapeutic strategies. Notably, the ability of D-NEMD, to detect subtle, mutation-dependent differences in early and long-range communication underscores the method’s sensitivity to functionally meaningful dynamical changes. These findings highlight the potential of D-NEMD to reveal how proteins reorganise their internal communication networks in response to sequence variation, opening new avenues for understanding, and ultimately targeting, allosteric regulation in complex biomolecular systems.

## METHODS

### Equilibrium MD simulations

Extensive equilibrium MD simulations (five replicates of 500 ns each) were carried out for each of the four systems comprising one EGFR kinase monomer bound at the ATP site by the non-covalent inhibitor BLU-945. All systems were constructed using the X-ray structure PDB ID 8D76,^41^ which contains three substitutions, namely T790M, L858R, and V948R. From this template, the following kinase-inhibitor complexes were generated: (*i*) a wild-type-like system in which L858R and T790M were reverted to their native residues while retaining V948R (labelled *wild-type*); (*ii*) a mutant system containing L858R and V948R, with residue 790 restored to threonine (labelled *L858R*); (*iii*) a system carrying L858R, T790M, and V948R (labelled as *L858R+T790M*); and, (*iv*) a control system in which all three substitutions (L858R, T790M, and V948R) were reverted to their wild-type forms (labelled *control*). A full description of system construction and equilibrium MD protocols is provided in the Supporting Information. This includes details on model preparation, protonation state assignment, energy minimisation, and initialisation procedures, simulation parameters, and all equilibrium analyses performed.

### D-NEMD simulations

A brief overview of the D-NEMD simulations is provided here; a full description of the methodology, simulation parameters, and analyses performed can be found in the Supporting Information.

The D-NEMD approach, originally proposed by Ciccotti *et al.*,^30,32^ combines simulations in equilibrium and nonequilibrium conditions to compute the dynamical behaviour of a system to an external perturbation (Fig. S12).^31^ This external trigger initially drives the system out of equilibrium, creating the driving force for the structural changes of interest to occur as the system relaxes towards a new equilibrium. Here, 450 short, 5 ns, nonequilibrium simulations were performed for the *wild-type*, L858R, *L858R+T790M,* and control systems to map the proteins’ response to removal of BLU-945 from the ATP binding site. Additionally, to account for the intrinsic motions of the protein, new equilibrium simulations (hereafter referred to as “null-perturbation” simulations) were initiated from the same 450 equilibrium conformations used for the nonequilibrium runs, after reassigning atomic velocities according to a Boltzmann distribution as described previously.^35,44^

The evolution of the proteins’ response to BLU-945 removal was then extracted using the Kubo-Onsager relation.^31,32^ The responses, described by the Cα displacement vector for each pair of nonequilibrium and equilibrium simulations, were computed after removing the protein’s equilibrium motions via the “null perturbation” analysis,^31,34^ with the convergence of Cα displacements assessed via the standard error of the mean (Fig. S12).

In this work, the instantaneous removal of the inhibitor from the ATP binding site of EGFR kinase domain and the analysis of how it adapts to an empty ATP binding site reveal the mechanical and dynamical coupling between this site and other structural elements, revealing the internal communication pathways within the system (Figs. S15-18).

## RESOURCE AVAILABILITY

### Lead contact

Requests for further information and resources should be directed to the lead contact, A. Sofia F. Oliveira.

### Data and code availability

All equilibrium and D-NEMD simulation data, including input files and trajectory files, will be made openly accessible via the University of Bristol data repository (data.bris) upon manuscript acceptance.

## ACKNOWLEDGMENTS

ASFO was supported at the University of Bristol by the Biological and Biotechnological Sciences Research Council ([BB/X009831/1]). ASFO and AJM thank the Biotechnology and Biological Sciences Research Council (BBSRC grant number BB/W003449/1). This work also received funding from the European Research Council (ERC) under the European Union’s Horizon 2020 research and innovation programme (Grant agreement No. 101021207; project information: PREDACTED). LT was funded at the University of Bristol by the “Heather Corrie Impact Fund”. MD simulations were carried out using the UK national supercomputer ARCHER2 and the GW4 Alliance Isambard-3 supercomputer. ASFO and AJM thank EPSRC via HECBIOSIM (hecbiosim.ac.uk) for providing ARCHER2 time.

## AUTHOR CONTRIBUTIONS

Conceptualisation-LT, ASFO and AJM; methodology-LT and ASFO; investigation-LT and ASFO; writing-original draft, LT, ASFO, AL and AJM; writing, review & editing-LT, ASFO, AL and AJM; funding acquisition-ASFO and AJM; supervision-ASFO and AJM.

