## Supporting Information for "Cancer-associated mutations reconfigure dynamical responses in EGFR kinase"

### Materials and methods

#### Model for *wild-type* and mutant EGFR kinases

A model for the intracellular kinase domain of EGFR was created based on the crystal structure 8D76.[1,2] This structure contains three substitutions, two of the most clinically impactful mutations, T790M and L858R, as well as V948R, a crystallisation-facilitating mutation commonly introduced to enhance protein stability during structure determination.[3] In structure 8D76, a non-covalent ligand, BLU-945, is bound in the ATP-binding site of the kinase domain.[1,2] BLU-945 (also known as tigozertinib or N-(2-((3S,4R)-3-fluoro-4-methoxypiperidin-1-yl)pyrimidin-4-yl)-5-isopropyl-8-((2R,3S)-2-methyl-3-(methylsulfonyl)methyl)azetidin-1-yl)isoquinolin-3-amine) is a fourth-generation, highly selective EGFR inhibitor developed by Blueprint Medicines to overcome resistance in NSCLC. Currently in phase 1/2 clinical trials (SYMPHONY Clinical Trial), BLU-945 exhibits sub-nanomolar potency against mutant EGFR (e.g. L858R/T790M: $IC_{50} \approx 0.4$ nM) and demonstrates >900-fold selectivity for the mutants compared to the wild-type kinase. [1,2] This structure corresponds to the active form of the kinase, characterised by the αC-helix adopting a closed conformation in which the stabilising K745–E762 salt bridge is intact, and the DFG motif positioned in the canonical “DFG-in” orientation.[4]

Given that the crystallisation-facilitating V948R substitution has been experimentally shown not to affect substrate affinity or catalytic activity,[3] this mutation was retained in all constructed models to preserve the structural environment of the crystallographic template as closely as possible. Using PDB ID 8D76 as the starting structure, we constructed three EGFR kinase–BLU-945 complexes: *(i)* a wild-type-like system in which L858R and T790M were reverted to their native residues while retaining V948R (hereafter referred to as *wild-type*); *(ii)* a mutant system containing L858R and V948R, with residue 790 restored to threonine (*L858R*); and *(iii)* a mutant system harbouring L858R, T790M, and V948R (*L858R+T790M*). To confirm that V948R does not influence the intrinsic functional dynamics of the kinase, we also constructed a fourth system (referred to as *control*) in which all three substitutions (L858R, T790M, and V948R) were reverted to their wild-type identities. Residue substitutions were performed using the PyMOL mutagenesis module[5] and selecting the residue orientation with fewer clashes.

The protonation states of titratable residues were assigned based on $pK_a$ values predicted by PROPKA,[6] and all systems were subsequently constructed using GROMACS.[7]

### Equilibrium molecular dynamics (MD) simulations of *wild-type* and mutant EGFR kinase

We collected five 500 ns-long MD simulations per system. The four models above were placed in a cubic box, solvated with TIP3P water[8] and 150 mM NaCl to resemble physiological concentration using GROMACS.[7] The AMBER99SB-ILDN force field[9] was used to describe the proteins, and BLU-945 was parametrized using ACPYPE[10] according to the General AMBER Force Field (GAFF).[11]
The systems were then minimized and heated following a three-step protocol using GROMACS.[7] A steepest descent algorithm was used during minimisation, with a 0.01 step size and considering a maximum of 500000 steps. From the minimised conformations, the systems were progressively heated in the NVT ensemble for 2 ns, reaching the temperature of 310 K using the first-order stochastic velocity rescaling thermostat and a temperature coupling time constant of 0.1 ps.[12] Five repeats were performed for each system, with their initial atomic velocities taken from a Boltzmann distribution. Following the NVT step, a 2 ns NPT simulation was performed, using the same thermostat as before. In the first two initialisation steps, restraints were applied on all heavy atoms using a force constant of 1000 kJ $mol^{-1}$ $nm^{-2}$. The third and final initialisation step, included a 2 ns simulation in the NPT ensemble, restraining only the Cα atoms in the protein (force constant of 1000 kJ $mol^{-1}$ $nm^{-2}$). Both NPT relaxation steps used the C-rescale barostat,[13] ensuring an exponential relaxation pressure coupling, including a stochastic term to enforce correct volume fluctuations.
All unrestrained simulations started from the last conformation of the third initialisation step. A 2 fs time step was used for the integration of Newton's equation of motion and the Parrinello-Rahman barostat[14] and V-rescale thermostat,[12] with a temperature and pressure time coupling constant of 0.1 ps and 0.1 ps, respectively. Non-bonded long-range electrostatic interactions were calculated using the smooth particle mesh Ewald method,[15] with a Fourier grid spacing of 1.6 Å and a 12 Å cut-off for direct contributions. A 12 Å cut-off was also used for the van der Waals interactions with long-range dispersion corrections for the energy and pressure. The LINCS algorithm[16] was used to constrain hydrogen-containing bonds in the protein and BLUE-945, and SETTLE[17] was used to keep water molecules rigid.

### Dynamical-nonequilibrium MD (D-NEMD) simulations

The D-NEMD approach, originally proposed by Ciccotti *et al.*,[18,19] combines simulations in equilibrium and nonequilibrium conditions (Fig. S12) to compute the dynamical response of a system to an external trigger, through the Kubo-Onsager relation.[20]

The rationale behind D-NEMD is simple and can be summarised as follows: when an external perturbation is applied to a system in equilibrium, a nonequilibrium trajectory can be initiated from the same configuration (Fig. S12), and the system's response extracted using the Kubo-Onsager relation by comparing the nonequilibrium and equilibrium trajectories at equivalent time points.[21,22] Despite its conceptual simplicity, D-NEMD provides a direct and powerful way of computing the system's dynamical response to external perturbations. A key advantage of D-NEMD is that the convergence and statistical significance of the responses can be rigorously assessed, with associated uncertainties readily reduced by increasing the number of nonequilibrium trajectories, ensuring sufficiently robust and reliable sampling and meaningful dynamical behaviours. This approach has been previously validated across fundamentally different protein systems from soluble enzymes to integral membrane proteins, and using a variety of external perturbations, including ligand removal, introduction of mutations, chemical reactions, and pH changes (e.g. references[23,24,25,26] ).

Starting from five independent 500-ns equilibrium MD simulations for each system, snapshots were extracted every 5 ns from the equilibrated part of the trajectories (50 to 500 ns), yielding 450 conformations per system. For each of these frames, BLU-945 was removed from the ATP site, and a new 5 ns-long nonequilibrium simulation was initiated (Fig. S12). BLU-945 removal initially drives the system out of equilibrium, creating the driving force for the structural changes of interest to occur as the kinase relaxes towards a new equilibrium. Similar ligand-removal perturbations have previously been used to map internal communication pathways in a range of systems, including β-lactamases,[27] nicotinic acetylcholine receptors,[28] SARS-CoV-2 spike variants,[29] among others.

To distinguish nonequilibrium responses from the protein's intrinsic equilibrium motions, an additional set of equilibrium simulations (hereafter referred to as "null perturbation" simulations) was generated from the same 450 conformations by reassigning atomic velocities according to a Boltzmann distribution (Fig. S12), following the procedure described previously.[25,30] Because no external perturbation is applied in this set, the systems remain at equilibrium throughout, and any structural changes observed in the "null perturbation" trajectories reflect only the proteins' inherent equilibrium dynamics.

In total, 3600 simulations (450 nonequilibrium and 450 "null-perturbation" trajectories per system) were carried out to assess how structural changes initiated at the ATP-binding site propagate across the kinase domain and how mutations reshape the protein's functional dynamics. All nonequilibrium and "null perturbation" simulations used GROMACS,[7] employing the same settings and conditions as the long equilibrium simulations above.

The proteins' response to the perturbation was quantified using the Kubo-Onsager relation,[19] by computing the displacement vector of each Cα atom between the equilibrium and nonequilibrium trajectories at corresponding time points (Fig. S12). At each time point, the Cα atom displacement vector was averaged over the 450 replicas. The resulting average displacement vectors indicate the direction of the proteins' motions following the perturbation (Figs. 3-5), with the norm of these vectors providing the magnitude of the Cα responses (Figs. 2-5 and Figs. S15-18).[21,22] The average responses shown in Figs. 15-18 were computed after removing the protein's equilibrium motions *via* the "null perturbation" analysis.[20]

To assess the convergence and significance of the responses, the standard error of the mean (SEM) was calculated.[21] A response was considered statistically significant only if its magnitude exceeded the SEM. Responses that did not meet this criterion were not considered. The resulting average displacements or the average displacement vectors were visualised on the closest structure to the average structure using the modevectors.py[31] plugin in PyMOL.[5]

### Supplementary figures

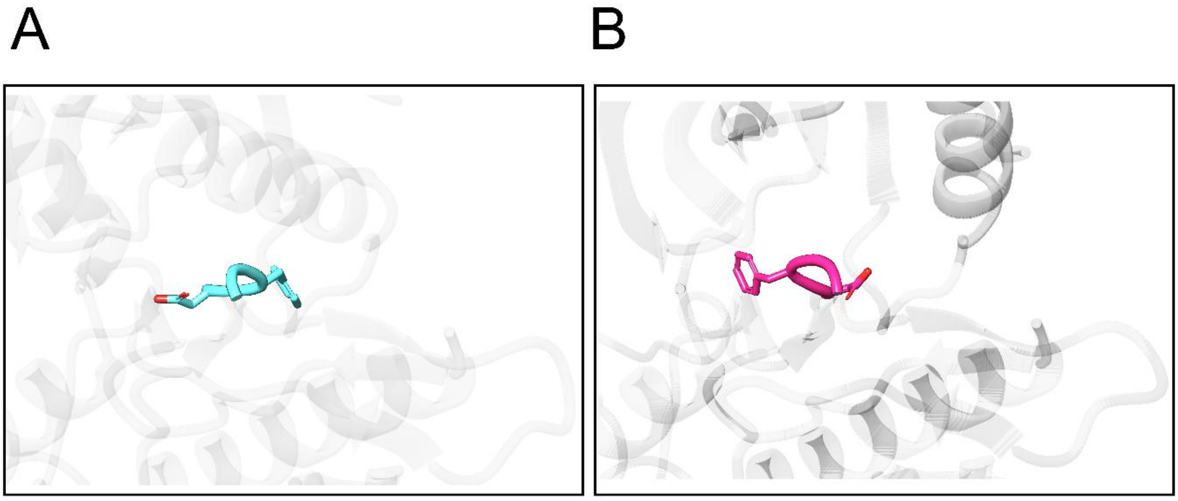


**Fig. S1: DFG-in vs DFG-out conformation.** The conformation of the DFG motif is a well-established marker of the kinase's activation state. **A)** Active DFG-in conformation (PDB ID:8D76).[1] **B)** Inactive DFG-out conformation (PDB ID:5HG5).[32] The protein is represented as a grey cartoon, and the residues D855, F856, and G857, which form the DFG motif, are shown as sticks.

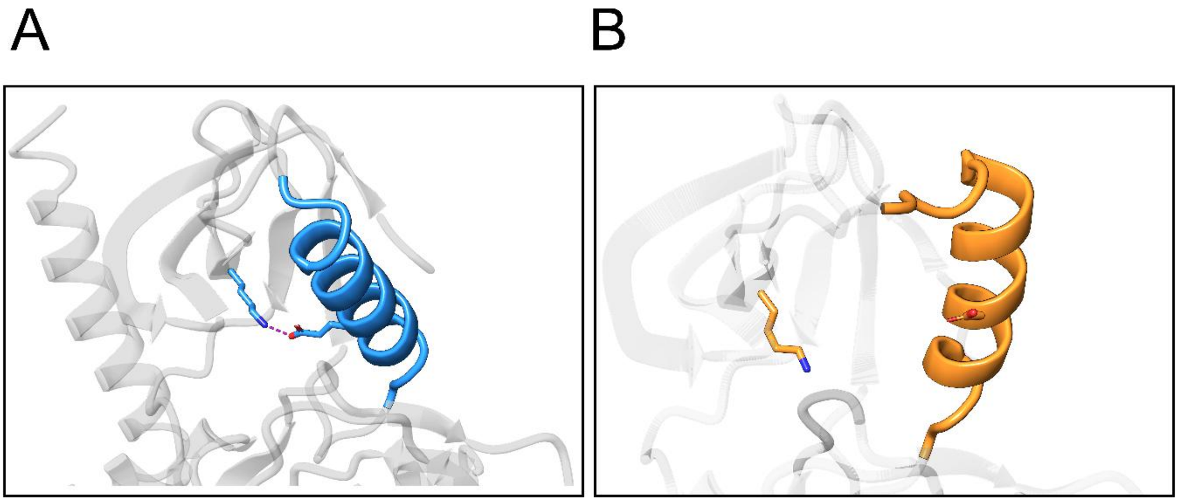


**Fig. S2: Comparison of active and inactive EGFR kinase domain conformation with focus on the αC-helix. (A)** Active-state conformation of the αC-helix, in which a stabilising salt bridge between K745 located next to the P-loop and E762 within the αC-helix is formed (PDB ID:8D76).[1] **(B)** Inactive-state conformation of the αC-helix, characterised by its outward rotation and loss of the K745–E762 salt bridge (PDB ID:5HG5).[32]

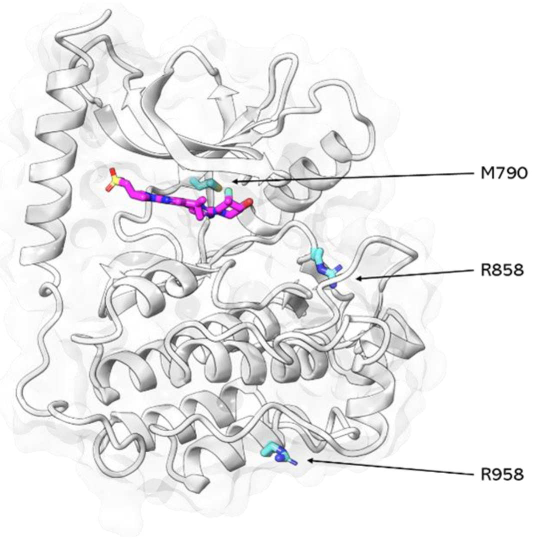


**Fig. S3: Crystal structure used as a starting point for this work.** The crystal structure of the L858R+T790M+V948R variant was used as a template for the equilibrium simulations (PDB ID:8D76). [1] The non-covalent inhibitor BLU-945, bound at the ATP-binding site, is shown with magenta sticks. BLU-945 is currently in phase 1/2 clinical trials for the treatment of resistant EGFR-driven NSCLC.[2] The cyan sticks highlight the position of the mutations.

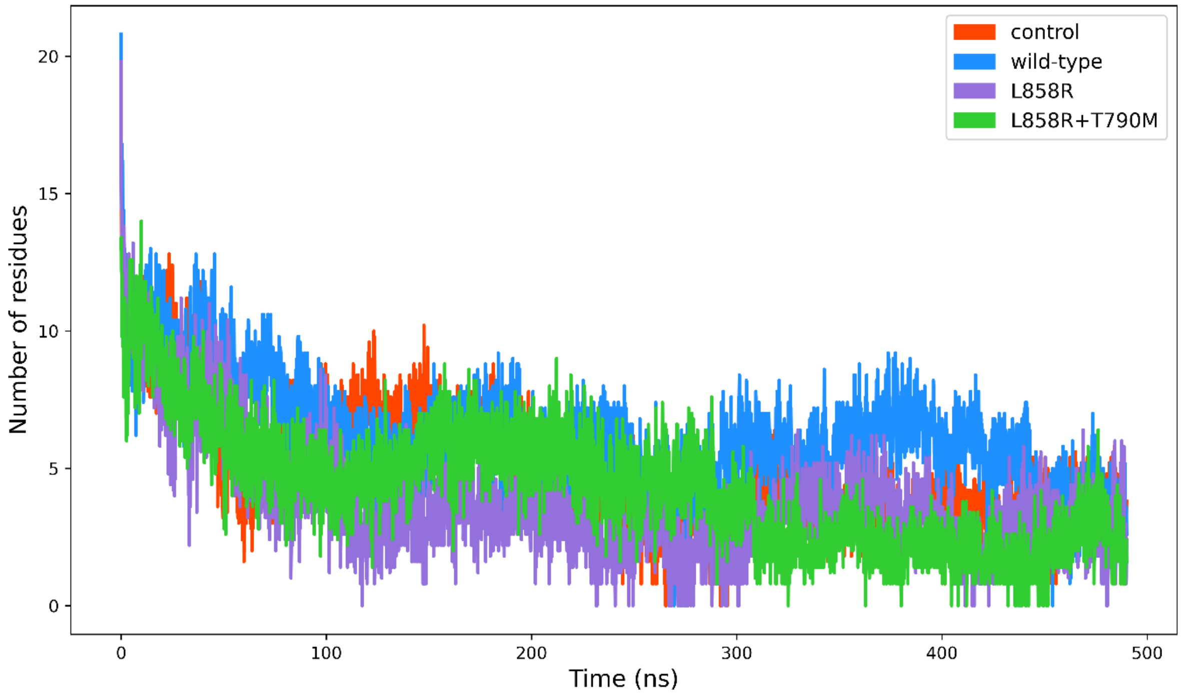


**Fig. S4: Time evolution of the secondary structure content of the tyrosine-rich region.** The average number of residues adopting helical secondary structure computed across all replicates for each system is shown. Secondary structure was assigned using DSSP,[33,34] considering all residues classified as α-helix, π-helix, $3_{10}$-helix, or 5-helix. Please note the loss of all helical content in the tyrosine-rich region after ~200 ns.

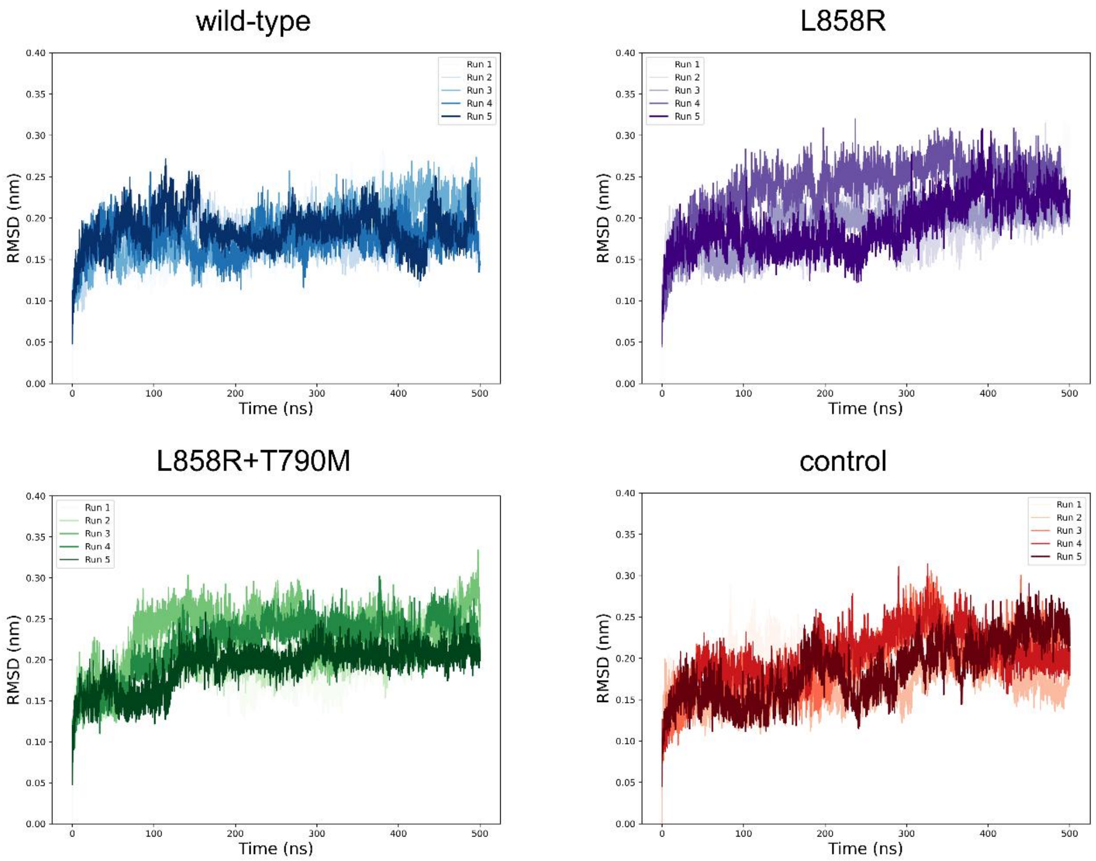


**Fig. S5: Temporal evolution of the Cα RMSD for wild-type and mutant systems.** The Cα RMSD was calculated relative to each system's starting structure and the averages obtained over all replicates for each system. Please zoom in on the image for detailed visualisation.

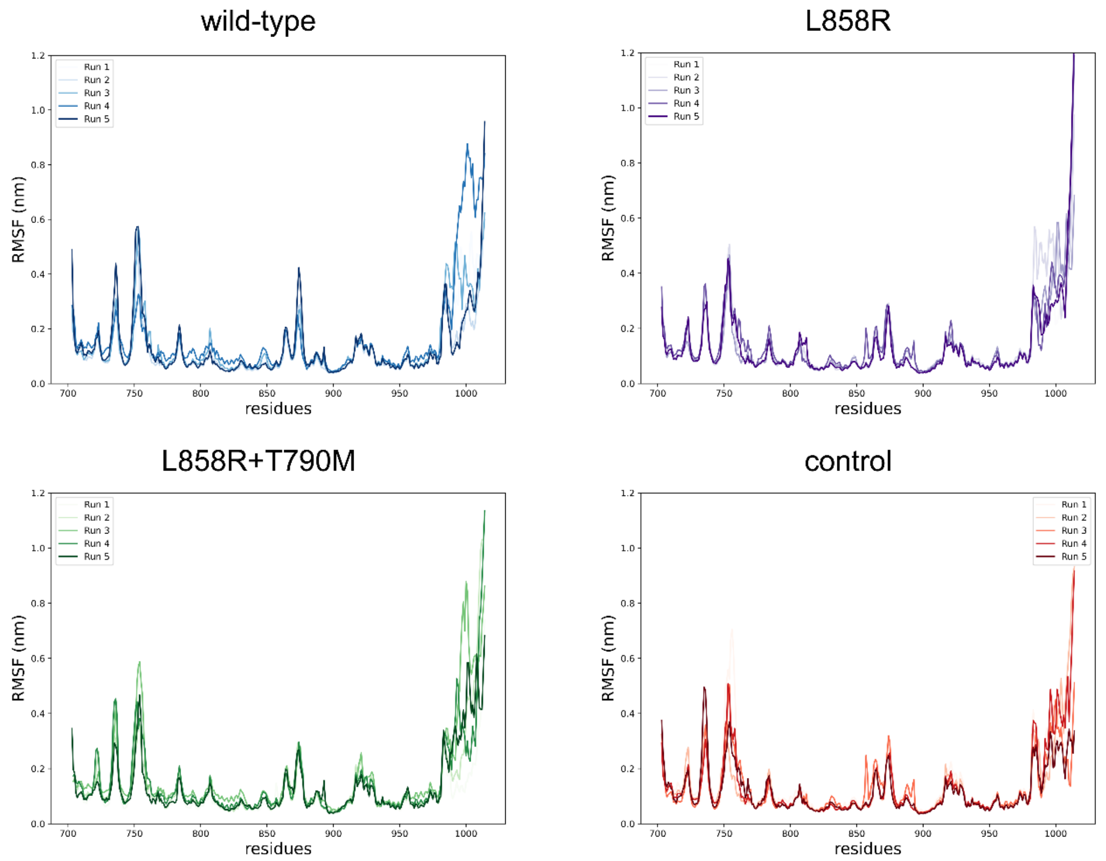


**Fig. S6: Cα RMSF for wild-type and mutant systems.** The Cα RMSF profiles were calculated using the equilibrated portion of the trajectories (50–500 ns) for each replicate. Please zoom in on the image for detailed visualisation.

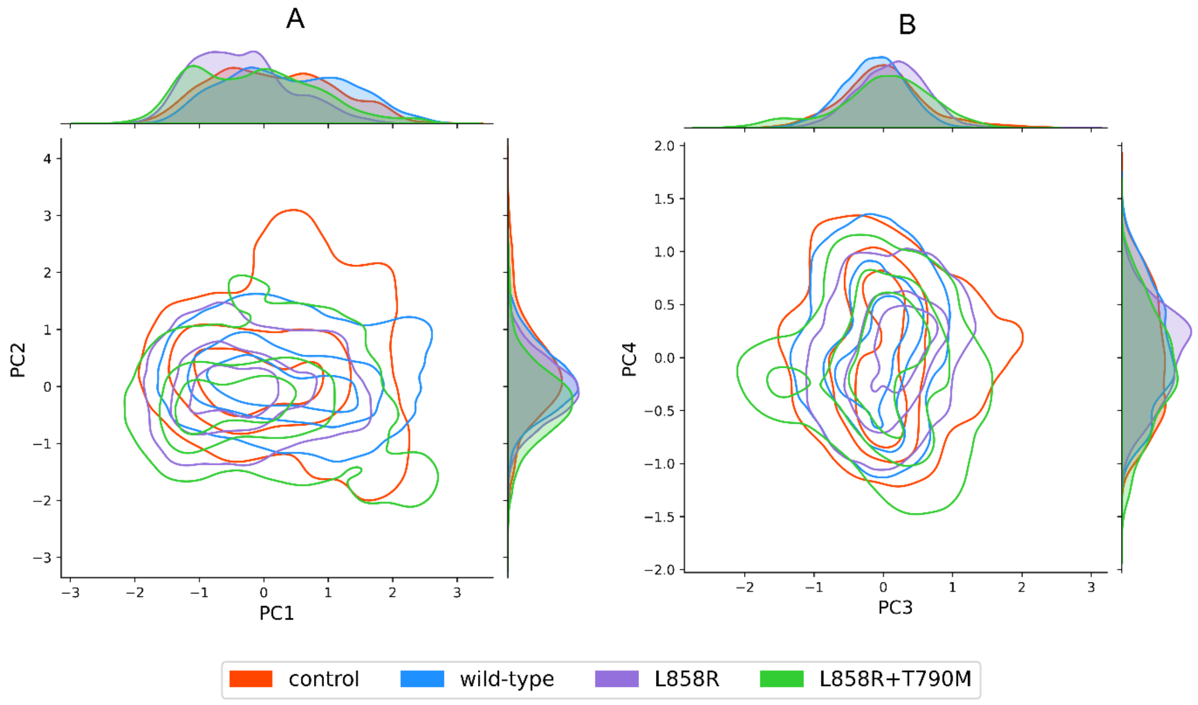


**Fig. S7: Principal component analysis (PCA) of the equilibrium simulations for each system.**

All replicates from all systems (namely *wild-type*, *L858R*, *L858R+T790M* and *control)* were combined into a single trajectory before analysis to ensure a shared conformational space, enabling direct comparison of the systems' behaviour. This trajectory contained one conformation every 100 ps per replicate and per system, with the proteins' Cα atoms, spanning 50-500 ns (totalling 90000 frames). Principal components (PC) 1-4 (PC1 vs PC2 on the left panel; PC3 vs PC4 on the right panel) were used to assess the extent to which the *wild-type*, *L858R*, *L858R+T790M* and *control* systems sample the conformational space. Overall, this analysis shows that all simulated systems occupy largely the same regions of the PC landscape, indicating broadly similar accessible conformational states.

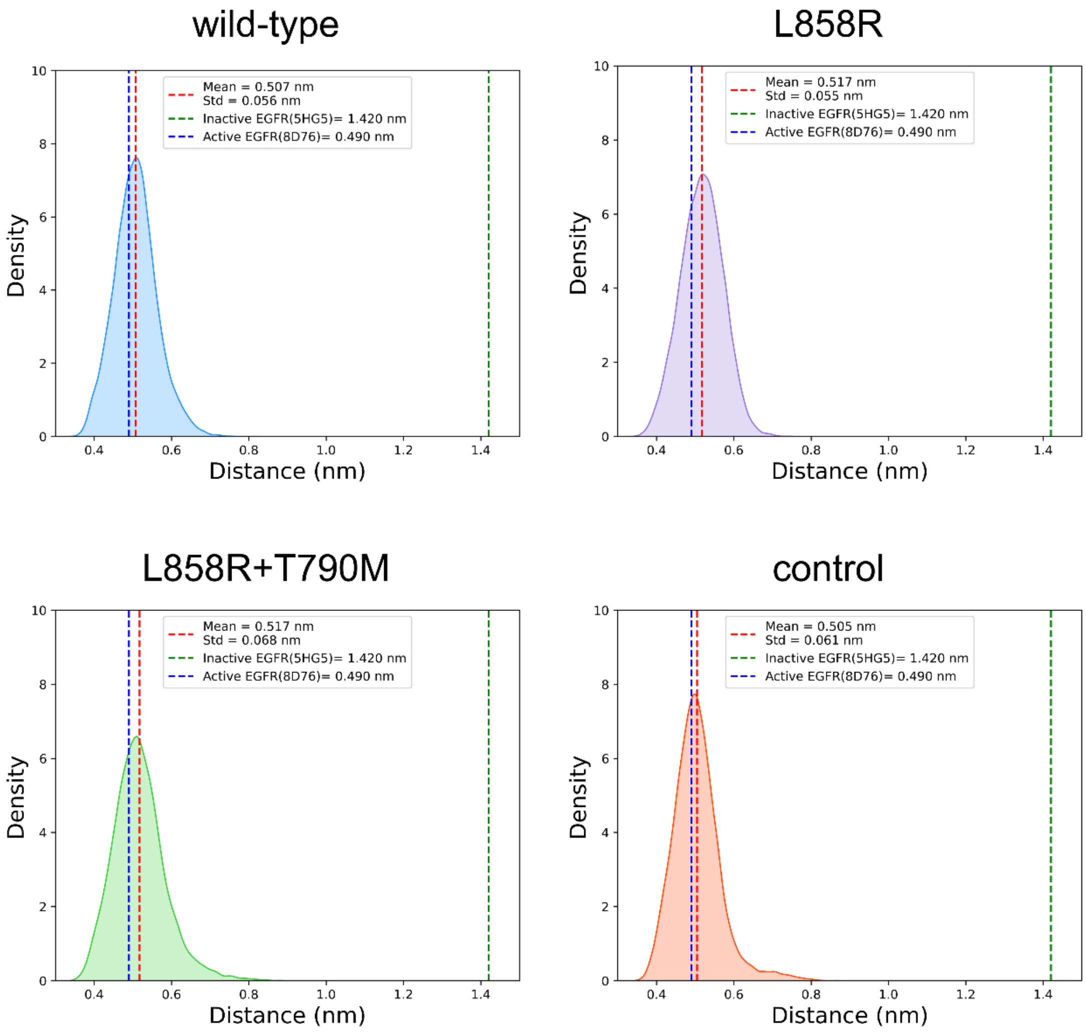


**Fig. S8: F856-M766 distance for the wild-type and mutant systems.** The distance between the centre of mass of the F856 side-chain and the Cα atom of M766 was calculated to assess whether the DFG motif adopted an active DFG-in or inactive DFG-out conformation during the equilibrium MD simulations. The histograms show the distribution of distances across all replicates for each system. Overall, the simulated complexes maintained the DFG-in conformation, indicating structural features characteristic of active EGFR. Please zoom in on the image for detailed visualisation.

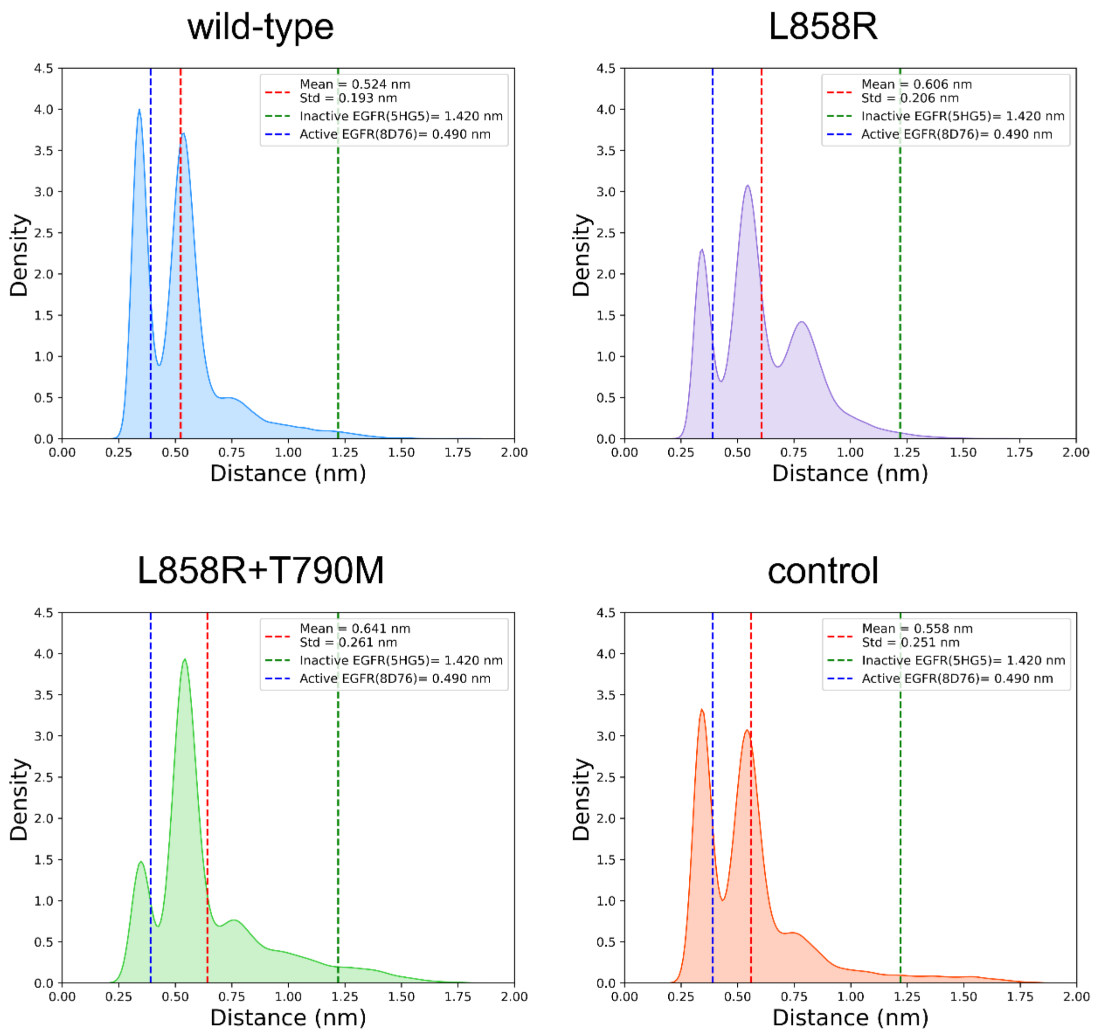


**Fig. S9: K745-E762 distance for the wild-type and mutant systems.** The distance between the nitrogen of K745 and the Cδ atom of E762 was calculated to assess whether the salt bridge between K745 and E762 (Fig.S2) was present during the simulations. The density plots show the distribution of distances across all replicates for each system. Overall, the simulated complexes maintained an average distance closer to the active form, resembling the semi-inactive form of the kinase described by Sutto *et al.*[35] Please zoom in on the image for detailed visualisation.

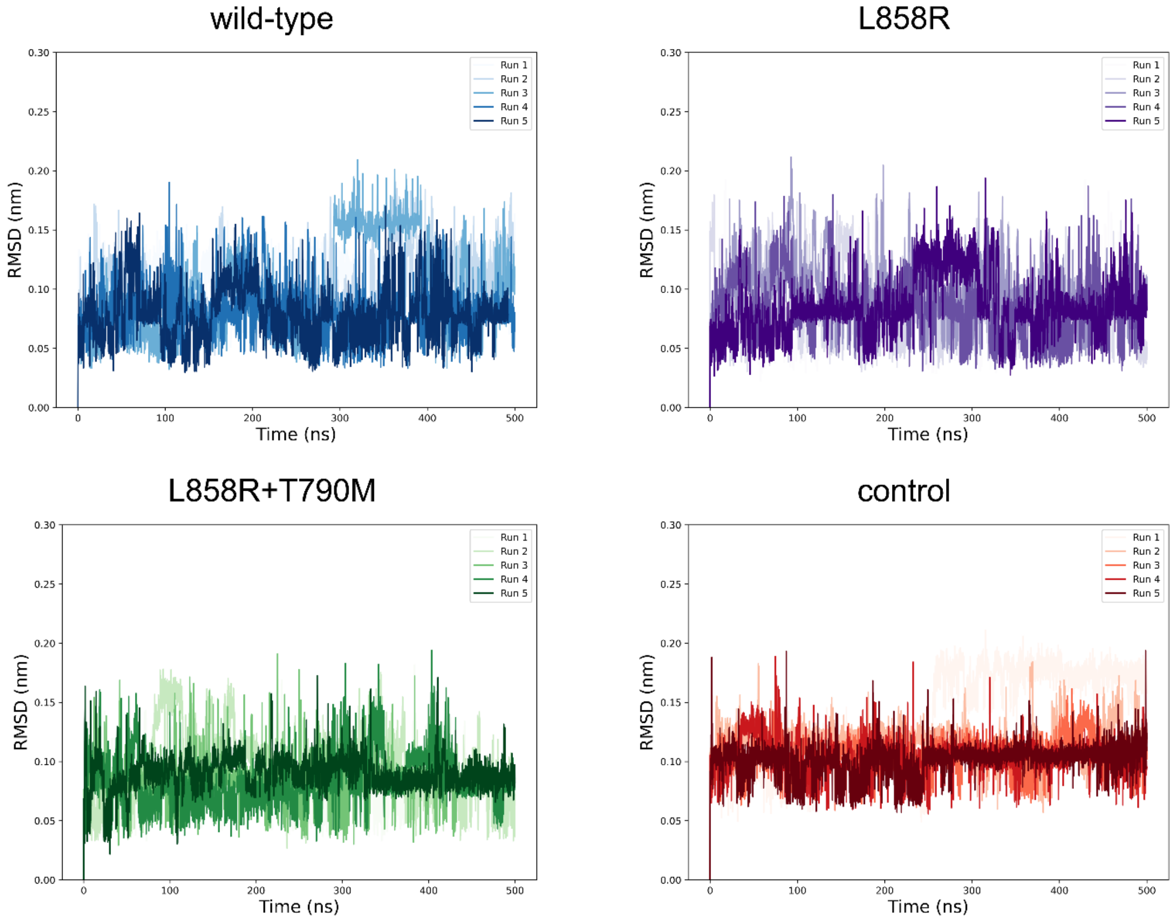


**Fig. S10: Temporal evolution of the RMSD for BLU-945 bound to the ATP site of the (A) *wild-type,* (B) *L858R*, (C) *L858R+T790M,* and (D) *control* systems.** The RMSD was determined relative to the initial binding mode of the inhibitor at the start of the simulations. This analysis shows that despite the changes in binding mode observed for the systems, BLU-945 remained stably bound to the protein throughout the simulation time.

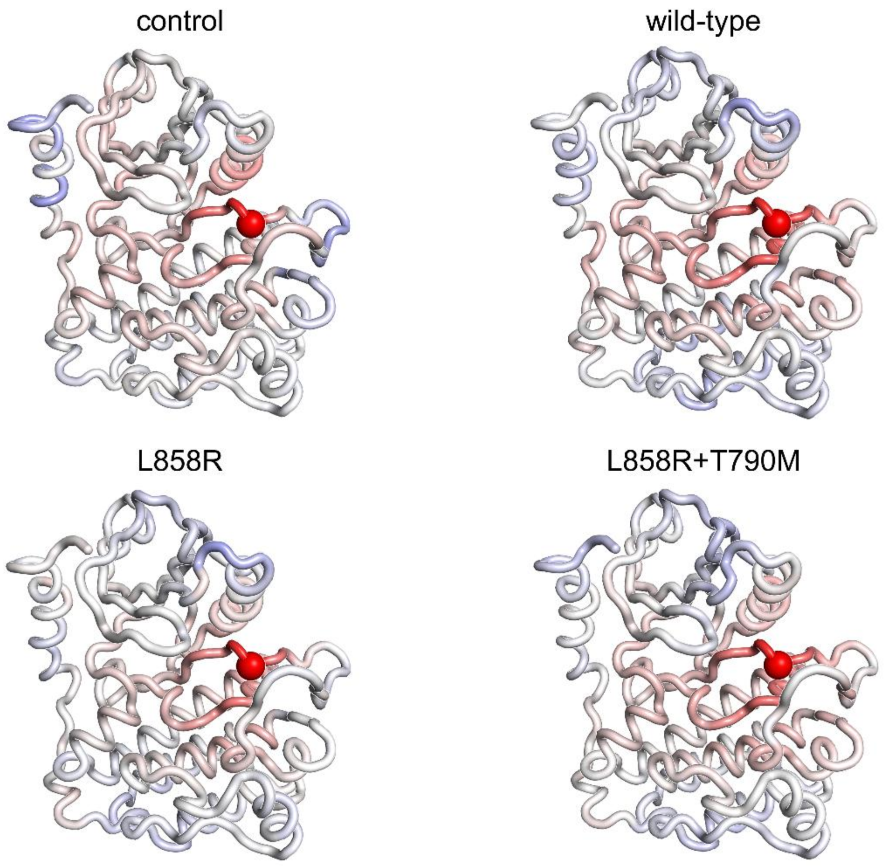


**Fig. S11: Statistical correlations for position 858 in the *wild-type*, *L858R*, *L858R+T790M,* and *control* systems.** The correlations between the Cα of 858 and all other Cα atoms in the kinase domain are shown. Note that the atoms that systematically move in opposite directions have a correlation value of -1, whereas those that systematically move in the same direction show a correlation of 1. The atoms whose movements relative to each other are uncoupled present a correlation value of 0. Fig. S11 shows that, in the equilibrium simulations, the mutations introduced do not drastically change the overall correlation between the Cα of residue 858 and the other Cα atoms in the kinase domain; therefore, the results obtained from the D-NEMD protocol can be mainly attributed to the applied perturbation (instantaneous removal of the ligand from the ATP-binding site).

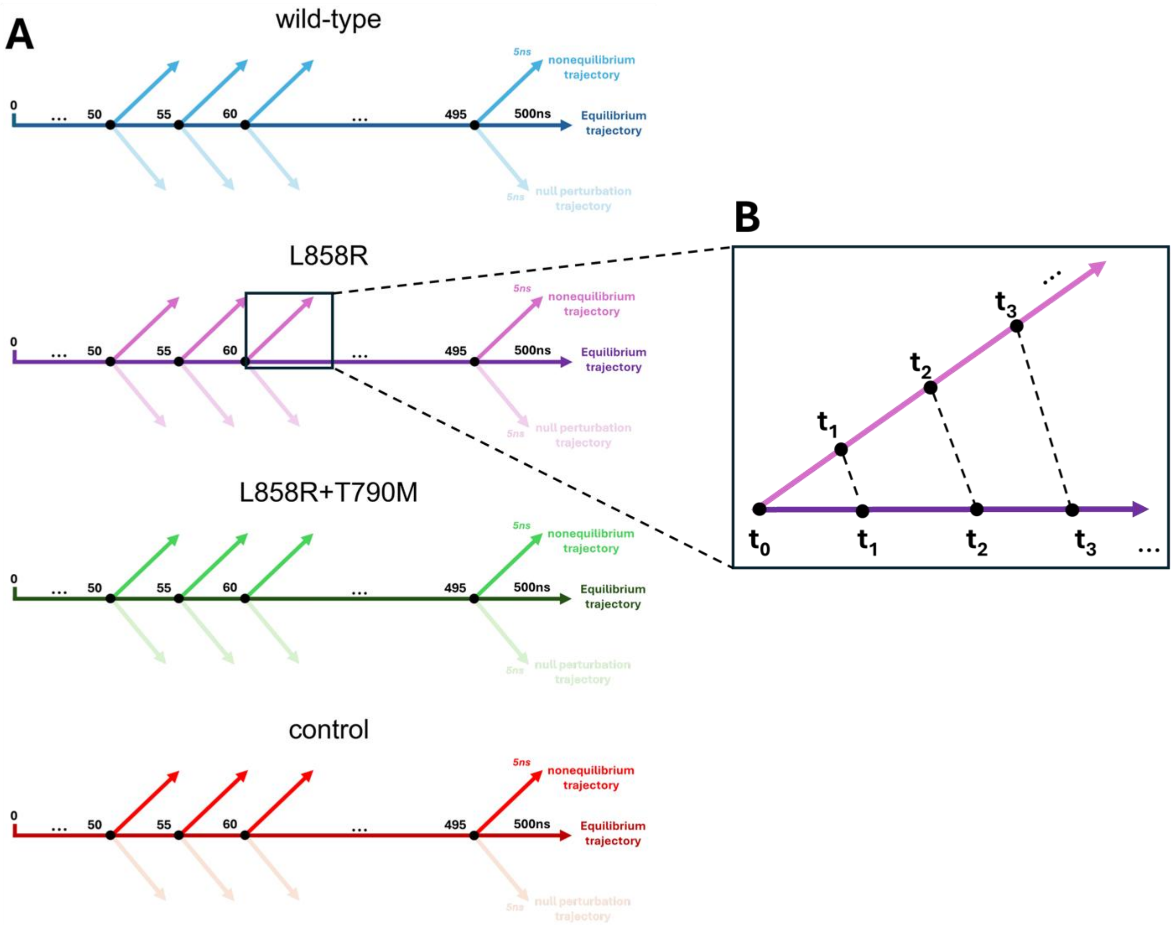


**Fig. S12: Schematic overview of the D-NEMD approach and analysis. (A)** Five equilibrium MD simulations, 500 ns each, were performed for the *wild-type*, *L858R*, *L858R+T790M,* and *control* systems with the non-covalent BLU-945 inhibitor bound at the ATP site. From the equilibrated part of each trajectory (from 50–500 ns), frames were extracted every 5 ns to serve as starting points for the short nonequilibrium simulations. Perturbations were applied by instantaneously deleting BLU-945 from the ATP-binding site, after which each nonequilibrium simulation was run for 5 ns. **(B)** The Kubo-Onsager relation[19,21] was used to extract the dynamical response of the system. For that, for each pair of equilibrium (with BLU-945 bound to the ATP site) and nonequilibrium (without BLU-945) trajectories, Cα displacement vectors were calculated at selected time points (0, 0.01, 0.1, 1, and 5 ns) and averaged across 450 simulations to capture the collective response. Zooming in is recommended for detailed visualisation.

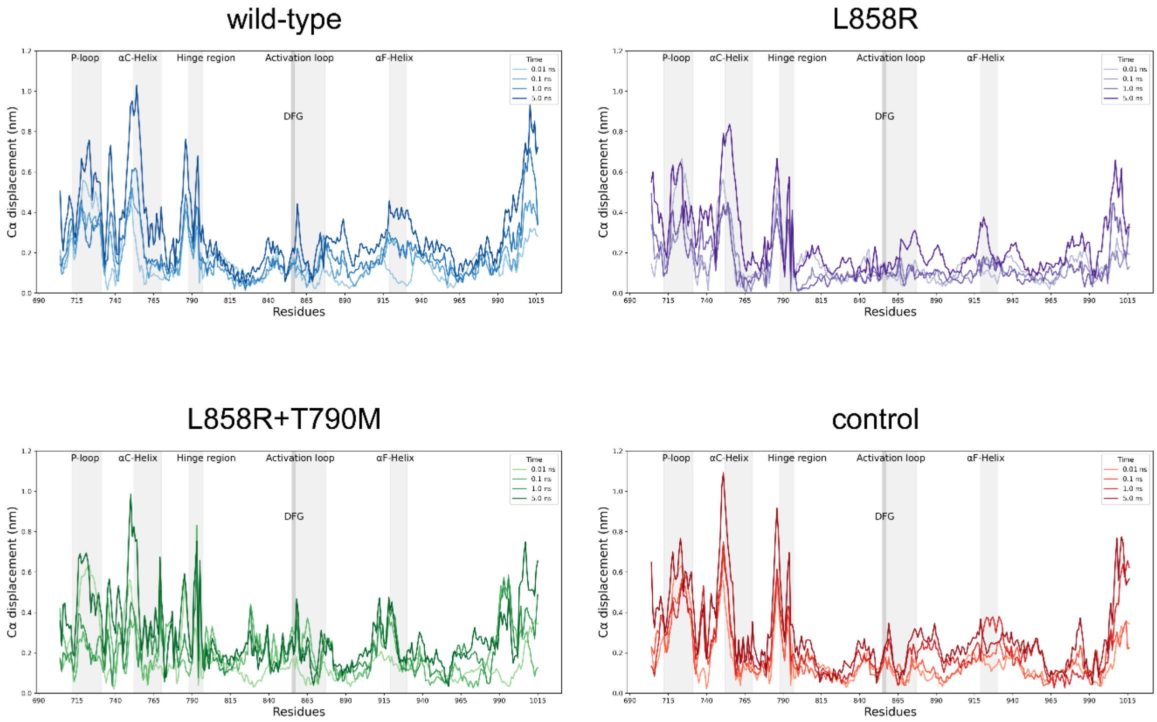


**Fig. S13: Dynamical responses of wild-type and mutant systems.** Amplitude of the Cα displacements at 0.1, 1, 2, and 5 ns following BLU-945 removal from the ATP site in: **(A)** *wild-type*, **(B)** *L858R*, **(C)** *L858R+T790M*, and **(D)** *control* systems. Average Cα displacement values were computed using the Kubo-Onsager relation[19,21] by comparing the nonequilibrium and equilibrium trajectories at equivalent times and averaging across all 450 replicates. The values shown correspond to the average responses after removing intrinsic protein fluctuations using the "null perturbation" analysis.[22,25] The positions of key structural motifs, including the P-loop, αC-helix, hinge region, activation loop, and DFG motif, are highlighted in grey. Zooming in is recommended for detailed visualisation.

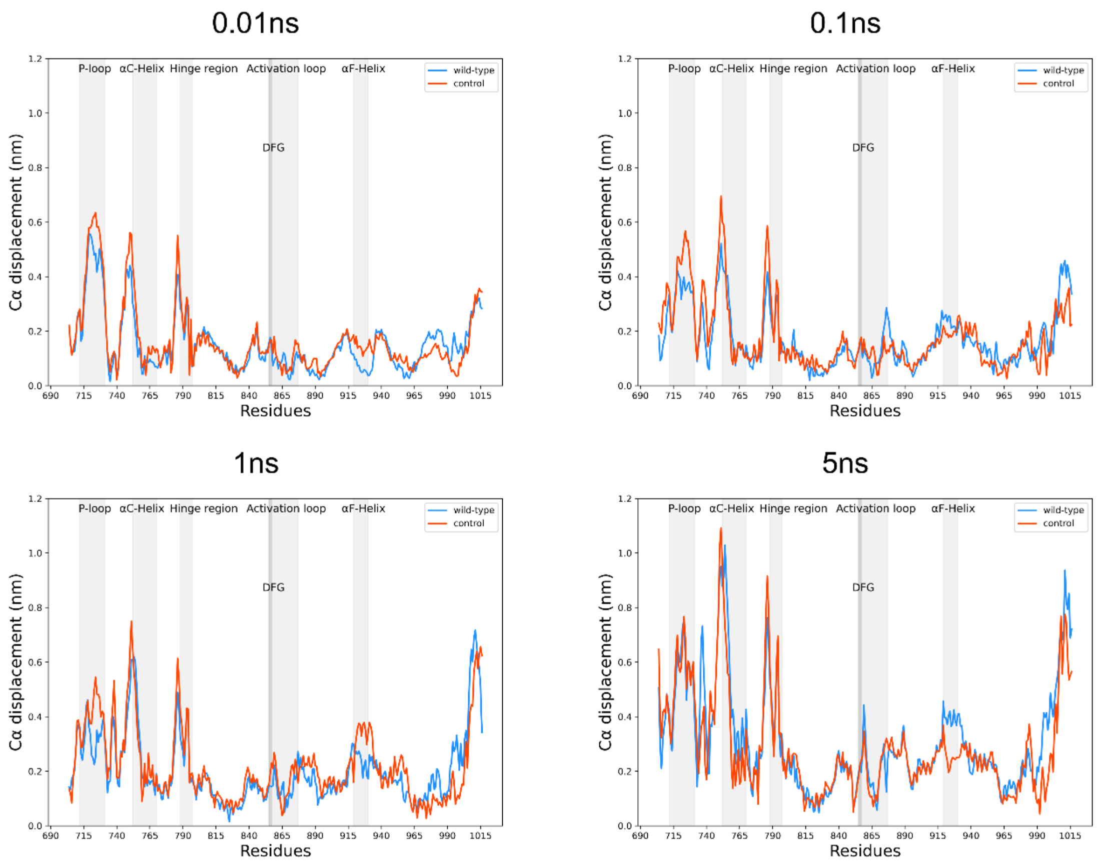


**Fig. S14: Dynamical responses of *wild-type* and *control* system.** Amplitude of the Cα displacements at 0.01, 0.1, 1, and 5 ns following BLU-945 removal from the ATP site in *wild-type (blue) and control (red)* systems. Average Cα displacement values were computed using the Kubo-Onsager relation[19,21] by comparing the nonequilibrium and equilibrium trajectories at equivalent times and averaging across all 450 replicates. The values shown correspond to the average responses after removing intrinsic protein fluctuations using the "null perturbation" analysis.[22,25]The position of key structural motifs, including the P-loop, αC-helix, hinge region, αF-helix, activation loop, and DFG motif, are highlighted in grey. As shown in the results, the two systems follow the same trend, confirming that the point mutation at position 948 (R948V) doesn't affect EGFR's dynamics. After 5 ns, there's a difference of slightly less than 0.2 ns only in the αF-helix; however, the directionality of the movement of this region is the same between the two systems. Zooming in is recommended for detailed visualisation.

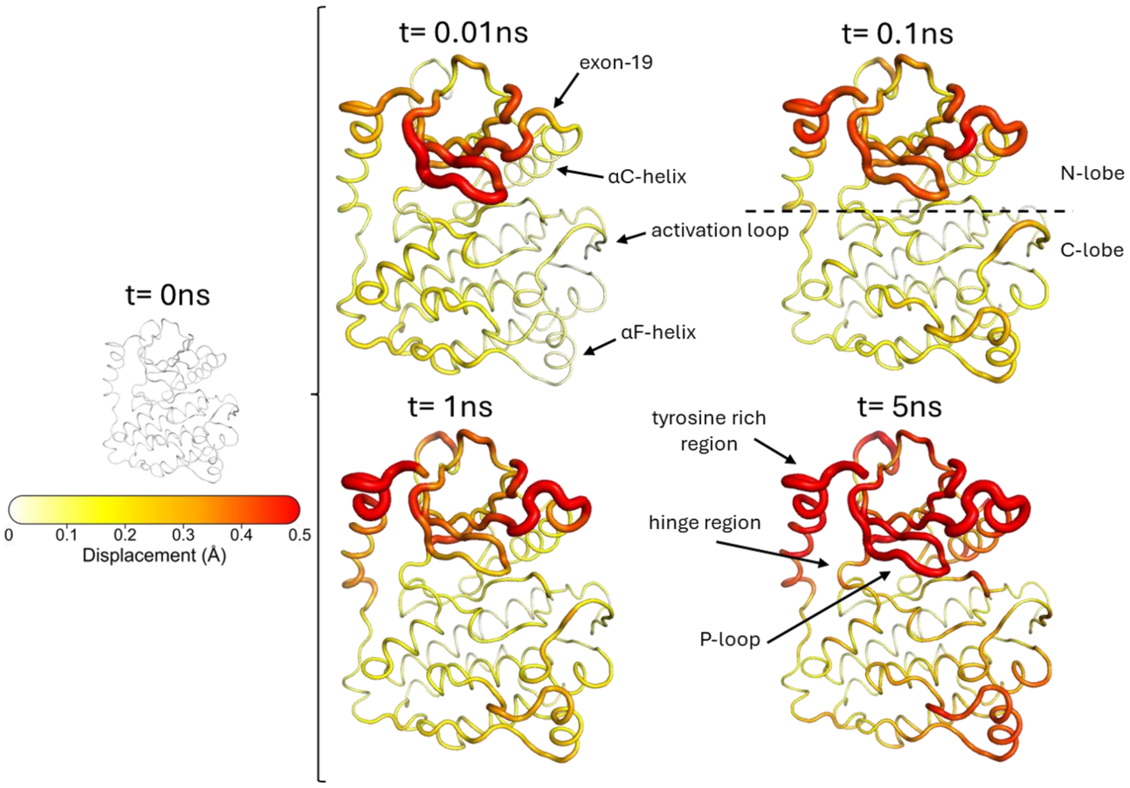


**Fig. S15: Dynamical response of the *wild-type* systems following the removal of BLU-945 from the ATP binding site.** The average Cα displacements after 0, 0.1, 1, and 5 ns are mapped onto the crystal structure 8D76.[1] The proteins' dynamical response was computed using the Kubo-Onsager relation, [19,21] by calculating the magnitude of the Cα displacement vectors between the equilibrium (with BLU-945 bound) and nonequilibrium (without BLU-945) trajectories at specific points in time for the *wild-type* system. The final displacement values correspond to the averaged D-NEMD responses obtained from all 450 trajectory pairs for each system, after removing the proteins' equilibrium fluctuations via the "null perturbation" analysis. [22,25] In this figure, the cartoon thickness and structure colours (scale on the left) indicate the amplitude of the proteins' response to BLU-945 removal.

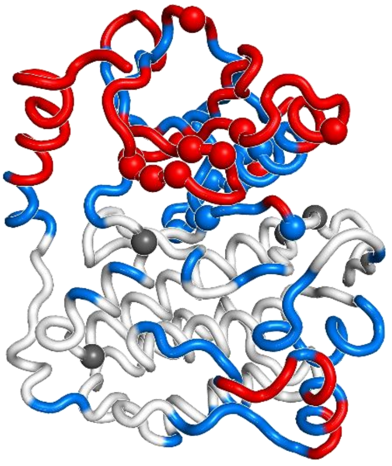

**Fig. S16: Location of substitutions, deletions and insertions reported for clinically relevant variants and their relationship to the allosteric pathways identified by D-NEMD.** The residues considered as hotspot for cancer-related mutations have been selected listing the positions studied by Robichaux et.al.[36]; namely, residues 709, 718, 719, 720, 724, 725, 747, 754, 763, 767, 768, 769, 770, 773, 774, 790, 792, 797, 811, 833, 854, 858, 861. The highest-responding regions identified by D-NEMD, namely the P-loop, αC-helix, hinge region, activation loop, and αF-helix, are coloured in red, where 61% of all the mutations taken into consideration fall directly into the identified regions. In blue are highlighted residues within second-shell contact distance (within 0.6 nm of the identified allosteric pathways), which, together with residues that fall directly within the D-NEMD pathways, account for 83% of the mutations. In grey, residues beyond the second-shell contact distance from the pathways identified by D-NEMD, which account for 17% of the mutations considered.

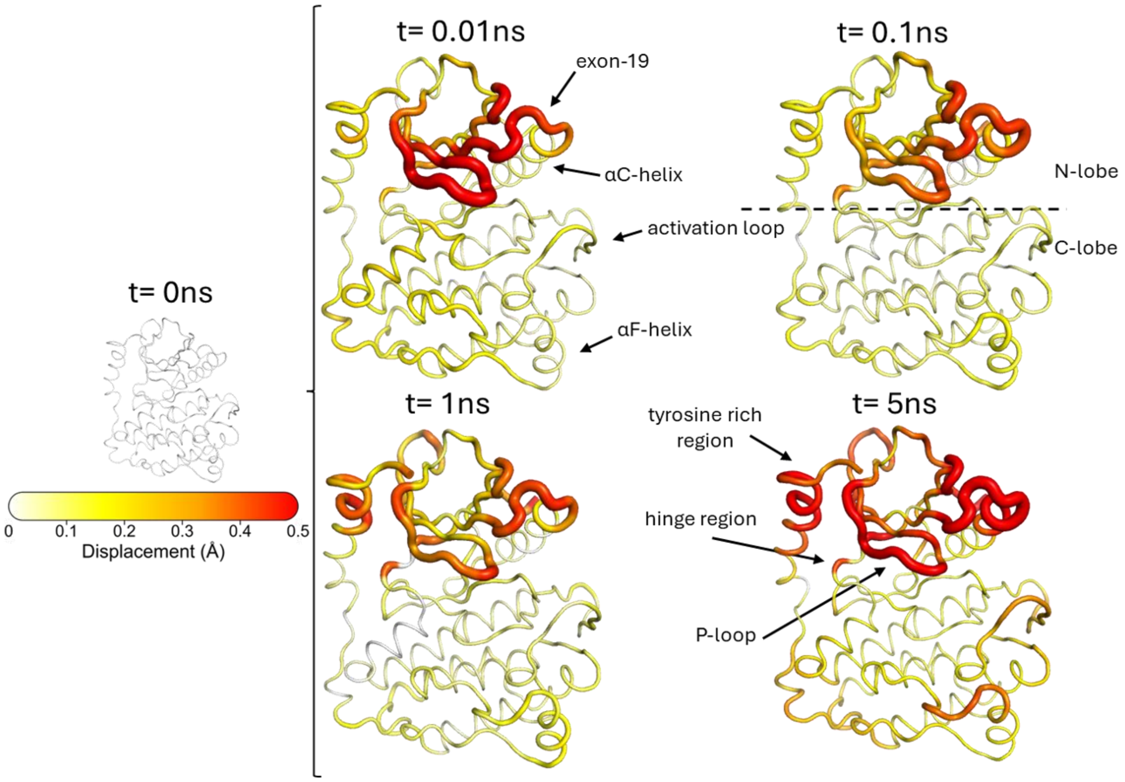


**Fig. S17: Dynamical response of the *L858R* systems following the removal of BLU-945 from the ATP binding site.** The average Cα displacements after 0, 0.1, 1, and 5 ns are mapped onto the crystal structure 8D76. [1] The proteins' dynamical response was computed using the Kubo-Onsager relation,[19,21] by calculating the magnitude of the Cα displacement vectors between the equilibrium (with BLU-945 bound) and nonequilibrium (without BLU-945) trajectories at specific points in time for the *L858R* system. The final displacement values correspond to the averaged D-NEMD responses obtained from all 450 trajectory pairs for each system, after removing the proteins' equilibrium fluctuations via the "null perturbation" analysis. [22,25] In this figure, the cartoon thickness and structure colours (scale on the left) indicate the amplitude of the proteins' response to BLU-945 removal.

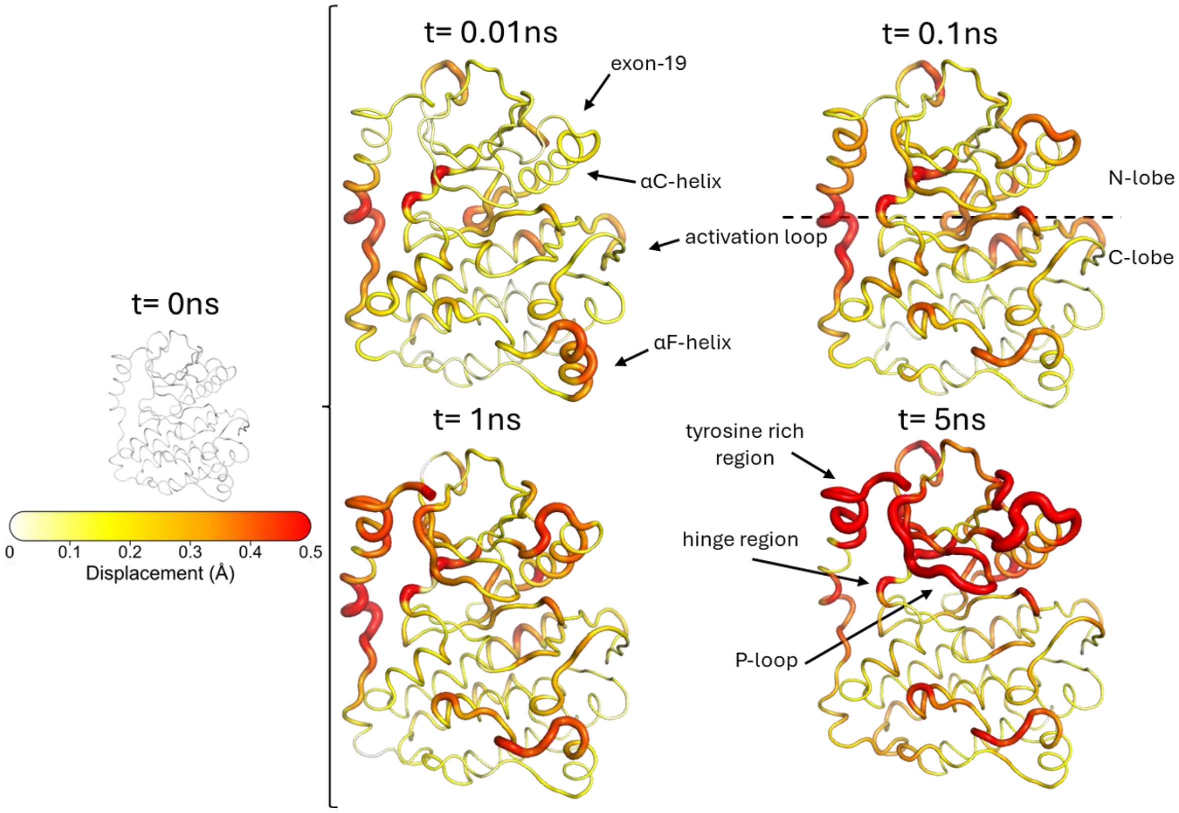


**Fig.S18: Dynamical response of the *L858R+T790M* systems following the removal of BLU-945 from the ATP binding site.** The average Cα displacements after 0, 0.1, 1, and 5 ns are mapped onto the crystal structure 8D76. [1] The proteins' dynamical response was computed using the Kubo-Onsager relation,[19,21] by calculating the magnitude of the Cα displacement vectors between the equilibrium (with BLU-945 bound) and nonequilibrium (without BLU-945) trajectories at specific points in time for the *L858R+T790M* system. The final displacement values correspond to the averaged D-NEMD responses obtained from all 450 trajectory pairs for each system, after removing the proteins' equilibrium fluctuations via the "null perturbation" analysis. [22,25] In this figure, the cartoon thickness and structure colours (scale on the left) indicate the amplitude of the proteins' response to BLU-945 removal.

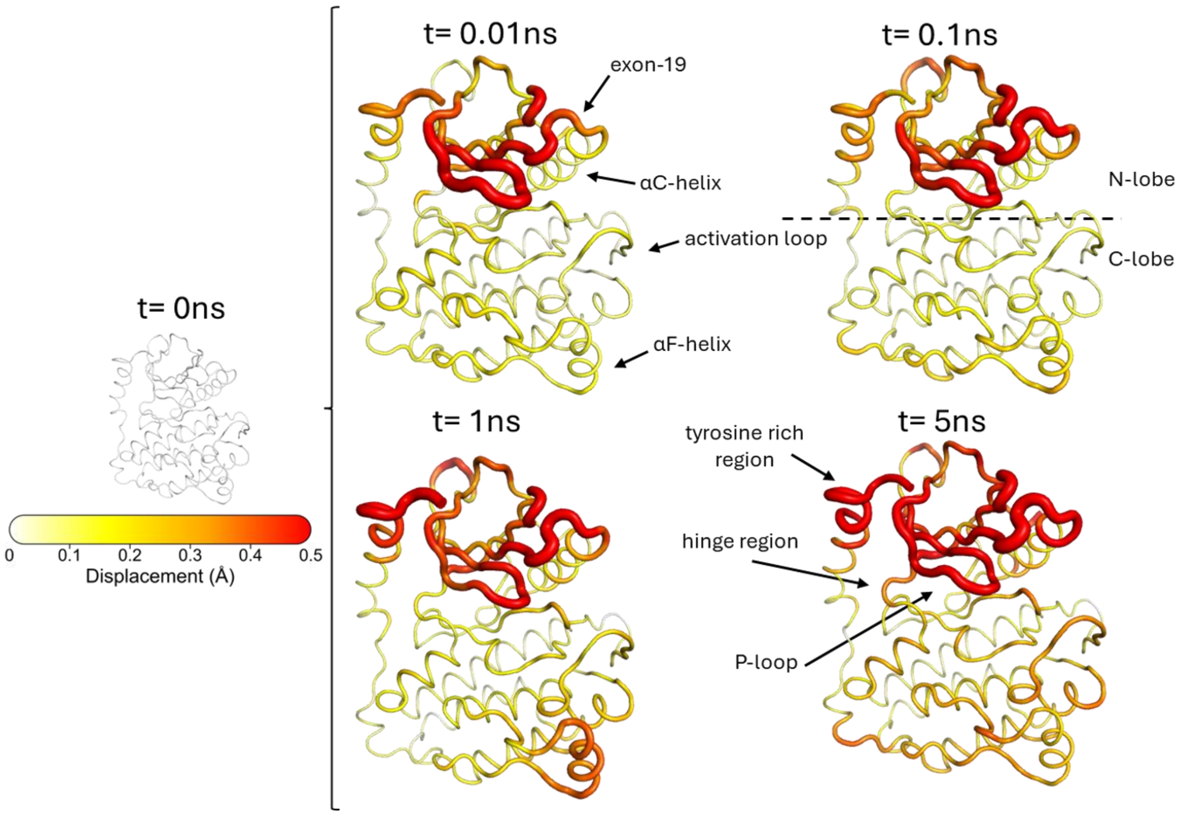


**Fig. S19: Dynamical response of the *control* systems following the removal of BLU-945 from the ATP binding site.** The average Cα displacements after 0, 0.1, 1, and 5 ns are mapped onto the crystal structure 8D76. [1] The proteins' dynamical response was computed using the Kubo-Onsager relation, [19,21] by calculating the magnitude of the Cα displacement vectors between the equilibrium (with BLU-945 bound) and nonequilibrium (without BLU-945) trajectories at specific points in time for the *control* system. The final displacement values correspond to the averaged D-NEMD responses obtained from all 450 trajectory pairs for each system, after removing the proteins' equilibrium fluctuations via the "null perturbation" analysis. [22,25] In this figure, the cartoon thickness and structure colours (scale on the left) indicate the amplitude of the proteins' response to BLU-945 removal.

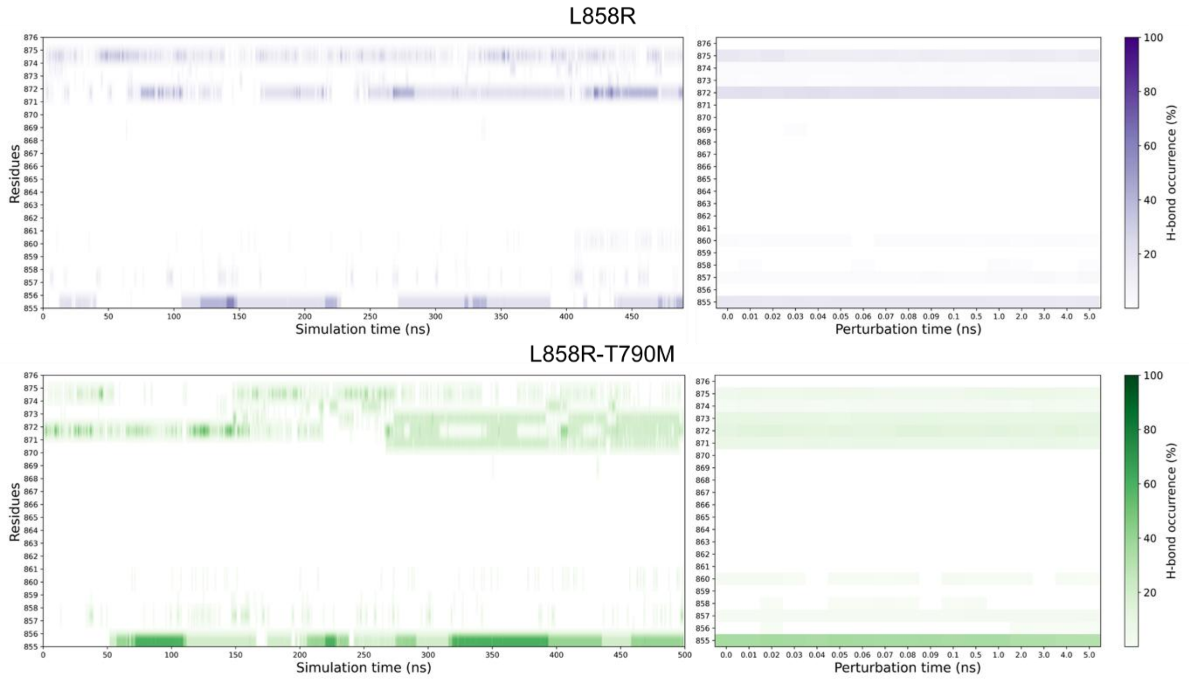

**Fig. S20: Hydrogen bond occurrence comparison between L858R and L858R+T790M with focus on the activation loop.** The frequency of forming a hydrogen bond between R858 and the residues within the activation loop (855-876) was calculated for *L858R* and *L858R+T790M* systems in both equilibrium and nonequilibrium simulations. This analysis evaluated the mutation's role in mediating interactions, particularly with the C-lobe. For the equilibrium simulations, hydrogen-bond occurrence was calculated as the percentage of frames in which the bond was present across five replicas at the same time point (one frame per replica). For the nonequilibrium simulations, the occurrence was calculated as the percentage of frames exhibiting the hydrogen bond across 450 trajectories at the same time point (one frame per nonequilibrium simulation).

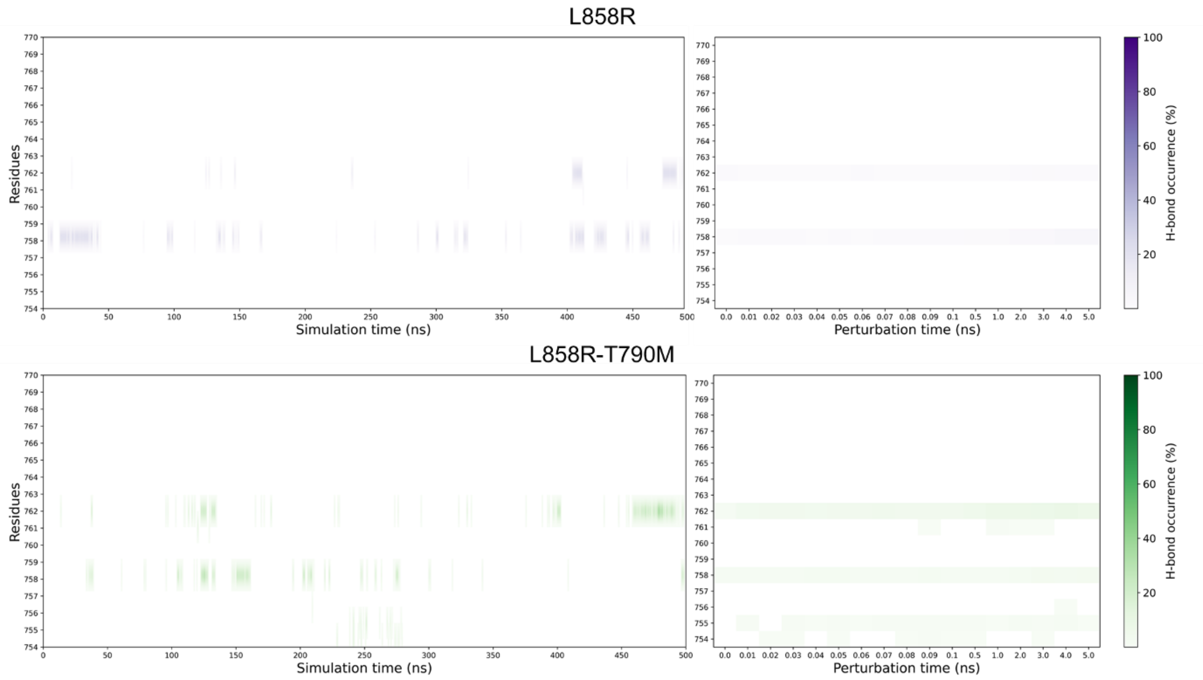


**Fig. S21: Hydrogen bond occurrence comparison between L858R and L858R+T790M with focus on αC-helix.** The frequency of forming a hydrogen bond between R858 and the residues within the αC-helix (754-770) was calculated for *L858R* and *L858R+T790M* systems in both equilibrium and nonequilibrium simulations. We performed this analysis to evaluate the mutation's role in mediating interactions, particularly with the αC-helix. For the equilibrium simulations, hydrogen-bond occurrence was calculated as the percentage of frames in which the bond was present across five replicas at the same time point (one frame per replica). For the nonequilibrium simulations, the occurrence was calculated as the percentage of frames exhibiting the hydrogen bond across 450 trajectories at the same time point (one frame per nonequilibrium simulation).
